# Microglial Neuropilin-1 trans-regulates oligodendrocyte expansion during development and remyelination

**DOI:** 10.1101/2021.01.07.425796

**Authors:** Amin Sherafat, Friederike Pfeiffer, Alexander M. Reiss, William M. Wood, Akiko Nishiyama

## Abstract

NG2 glia or oligodendrocyte precursor cells (OPCs) are distributed throughout the gray and white matter and generate myelinating cells. OPCs in white matter proliferate more than those in gray matter in response to platelet-derived growth factor AA (PDGF AA), despite similar levels of its alpha receptor (PDGFRα) on their surface. Here we show that the type 1 integral membrane protein Neuropilin-1 (Nrp1) is expressed not on OPCs but on amoeboid and activated microglia in white but not gray matter in an age- and activity-dependent manner. Microglia-specific deletion of Nrp1 compromised developmental OPC proliferation in white matter as well as OPC expansion and subsequent myelin repair after acute demyelination. Exogenous Nrp1 increased PDGF AA-induced OPC proliferation and PDGFRα phosphorylation on dissociated OPCs, most prominently in the presence of suboptimum concentrations of PDGF AA. These findings uncover a novel mechanism of regulating oligodendrocyte lineage cell density that involves trans-activation of PDGFRα on OPCs via Nrp1 expressed by adjacent microglia.

## INTRODUCTION

Oligodendrocyte precursor cells (OPCs), also known as NG2 glia or polydendrocytes, are widely distributed throughout the developing and mature central nervous system (CNS) ^1-3^, where they comprise 2-9% of the total cells. The developmental expansion of this population is critically dependent on platelet-derived growth factor AA (PDGF AA) acting on their alpha receptor (PDGFRα) ^4-6^. The most characterized and established role of OPCs is their ability to self-renew and generate the correct number of myelinating oligodendrocytes (OLs) needed for optimal network function. Hypomyelinating or dysmyelinating mouse mutants with reduced OLs or myelin are accompanied by increased OPC proliferation ^7,8^. In the mature CNS, a demyelinated lesion rapidly elicits OPC proliferation ^9-12^. However, the molecular and cellular mechanism by which such feedback signals promote OPC proliferation is not known.

While OPCs are evenly distributed throughout the CNS, the rate of OPC proliferation and OL differentiation is greater in white matter than in gray matter ^13,14^, and those in white matter proliferate more in response to PDGF AA ^15^. While both cell intrinsic and extrinsic factors influence the differential OPC behavior ^16^, heterotopic transplantation of 300-μm^3^ pieces in slice cultures suggests that the greater proliferative response of OPCs in white matter to PDGF AA is imparted by their local pericellular microenvironment ^15^.

To determine the mechanism underlying the differential response of OPCs in gray and white matter to PDGF AA, we searched the literature for a potential co-receptor for PDGF that could differentially regulate the proliferative response of OPCs to PDGF AA in gray and white matter. We identified heparan sulfate proteoglycans (HSPGs), and neuropilin-1 (Nrp1) ^17,18^ as potential candidates. Since heparin and HSPGs are known to affect a variety of growth factors, we chose to explore the role of Nrp1 as a molecule with more specific targets. Nrp1 is a type 1 transmembrane molecule, which was first discovered in the developing *Xenopus* optic tectum ^19^ and subsequently shown to be expressed on developing murine axons ^20^ and regulate axon pathfinding by binding to class III Semaphorins ^21^. In endothelial cells, Nrp1 binds vascular endothelial cell growth factor (VEGF) and modulates signal transduction through VEGF receptor (VEGFR) ^22^. Here we show that Nrp1 is expressed on amoeboid and activated microglia in the developing and demyelinated corpus callosum and promotes OPC proliferation by activating PDGFRα on OPCs *in trans*.

## METHODS

### Animals

All animal procedures were approved by the Institutional Animal Care and Use Committee of the University of Connecticut. For slice cultures, we used NG2cre;Z/EG double transgenic mice ^15^, which were obtained by breeding NG2cre homozygous mice to Z/EG homozygous or heterozygous mice ^23^ (Jackson Laboratory stock no. 003920, Tg(CAG-Bgeo/GFP)21Lbe/J; RRID:IMSR_JAX:003920). The NG2cre mice used in this study had been generated by injecting a BAC construct that contained NG2creER™, but one of the founders (NG2creERA) expressed cre independently of tamoxifen ^24^ and exhibited reporter expression in a temporal and spatial pattern that was identical to NG2creBAC;Z/EG mice previously described ^25^. Since cre in the NG2creERA line was more stable than that in the original NG2creBAC mice described in ^25^, the slice culture experiments described here were performed with NG2creERA;Z/EG mice, which we will refer to here as NG2cre;Z/EG mice. To delete Nrp1 in microglia, we crossed Nrp1^fl/+^ or Nrp1^fl/fl^ mice ^26^ (Jackson Laboratory stock no. 005247, B6.129(SJL)-*Nrp1*^*tm2Ddg*^/J; RRID:MGI:3528190) with Cx3cr1cre^ERT2-ires-EYFP^ mice ^27^ (Jackson Laboratory stock no. 021160B6.129P2(Cg)-*Cx3cr1*^*tm2*.*1(cre/ERT2)Litt*^/WganJ; RRID:MGI:5528845) to generate mg-Nrp1-cont or mg-Nrp1-cko mice, respectively. Both males and females were used in all the experiments. To induce cre in mg-Nrp1-cont or mg-Nrp1-cko mice for developmental studies, we injected 4-hydroxytamoxifen (4OHT, Sigma H7904, 100 μg/g) at P2 and P3 and used the mice at P8 for slice cultures or perfused them at P5, P8, P14, and P30 for histological analysis. To induce cre in weaned mice, 100 μg/g tamoxifen (Cayman Chemicals 13258 or Sigma T5648) was injected daily intraperitoneally for 4 consecutive days. To detect proliferating cells, 5-ethynyl-2’-deoxyuridine (EdU, 50 μg/g, Cayman Chemicals 20518) was injected intraperitoneally twice, 2 hours apart, prior to sacrifice.

#### Demyelinating lesion

To induce demyelination, we injected 2 μl of 2.5% lysolecithin (LPC, α-lysophosphatidylcholine, Sigma L4129) dissolved in PBS into the corpus callosum of 9-to 12-week-old mg-Nrp1-cont and mg-Nrp1-cko mice, using the stereotaxic coordinates 0.3 mm anterior from those previously described ^28^. The animals were perfused at 3, 7, 14, and 28 days post lesioning (dpl). To examine OPC proliferation, EdU was injected twice, 2 hours apart, before sacrifice at 3 or 7 dpl.

### Tissue processing and immunohistochemistry

Mice were perfused with 4% paraformaldehyde (PFA) containing 0.1M L-lysine and 0.01M sodium meta-periodate and postfixed in the same fixative for 2 hours, after which the tissues were dissected and washed 4 times in 0.2M sodium phosphate buffer, pH7.4. The tissues were cryoprotected in 0.2M sodium phosphate buffer containing 30% sucrose for at least 24 hours, frozen in OCT compound (Tissue-Tek; Adwin Scientific 14-373-65), and 20-μm sections were cut on a Leica CM3050S cryostat. For free floating sections, 50-μm-thick coronal sections were cut with a Leica vibratome VT1000S directly after fixation and rinsing.

For immunolabeling, sections were rinsed in PBS, blocked in 5% normal goat serum (NGS) or 1% bovine serum albumin (BSA) containing 0.1% Triton X-100 in PBS for 1 hour at room temperature, followed by incubation in the primary antibodies at 4°C overnight. The antibodies are listed in **Table 1**. The sections were rinsed 3 times in PBS and incubated in the secondary antibodies at room temperature for 1 hour. Then the sections were rinsed and mounted with Vectashield with DAPI (Vector, H-1200). To detect EdU, immunolabeled sections were washed 3 times with PBS and incubated in the Click reaction mixture containing 150 mM NaCl, 100 mM TrisHCl pH7.15, 4 mM CuSO_4_.5H_2_O (Sigma, C6283), 4 ng/mL Alexa Fluor-647-conjugated azide (ThermoFisher, A10277), and 100 mM sodium ascorbate (Sigma, A4034) at room temperature for 30 minutes. The sections were then washed 3 times in PBS, stained with 5 μg/mL Hoechst 33342 in PBS (InVitrogen Molecular Probes, LSH3570), and mounted in Vectashield without DAPI (Vector, H-1000). To examine cell death, we applied ApopTag Red In Situ Apoptosis Detection Kit (Millipore, S7165) according to the manufacturer’s recommended procedure, after treating the tissues with pre-cooled ethanol:acetic acid (2:1) for 5 minutes.

**Table 1.** Primary and secondary antibodies used

| Antibody | Source | dilution | cat no | RRID |
| --- | --- | --- | --- | --- |
| **Primary antibodies** |  |  |  |  |
| Mouse anti-CC1 | Millipore | 1:200 | OP80 | RRID:AB_2057371 |
| Rat anti-CD140a, clone APA5 | BD Sciences |  | 558774 | RRID:AB_397117 |
| Rat anti-CD68 | Biolegend | 1:1000 | 137001 | RRID:AB_2044003 |
| Rat anti-F4/80 | Bio-Rad | 1:200 | MCA497R | RRID:AB_323279 |
| Chicken anti-GFP | Aves Laboratories | 1:500 | GFP-1010 | RRID:AB_2307313 |
| Mouse anti-glyceraldehyde-3-phosphodehydrogenase (GAPDH) | Millipore | 1:2000 | MAB374 | RRID:AB_2107445 |
| Goat anti-Iba1 | Invitrogen | 1:500 | PA5-18039 | RRID:AB_10982846 |
| Rabbit anti-laminin | Sigma | 1:1000 | L8271 | RRID:AB_477162 |
| Mouse anti-myelin basic protein (MBP, SMI99) | Biolegend | 1:2000 | 808403 | RRID:AB_2734562 |
| Mouse anti-Neurofilament H (nonphosphorylated, SMI32; 1:3000) | Biolegend | 1:3000 | 801701 | RRID:AB_2564642 |
| Rabbit anti-NG2 | Millipore | 1:500 | AB5320 | RRID:AB_11213678 |
| Goat anti-Nrp1 | R&D Systems | 1:200 | AF566 | RRID:AB_355445 |
| Mouse anti-Olig2 | Millipore | 1:500 | MABN50 | RRID:AB_10807410 |
| Goat anti-PDGFRα | R&D Systems | 1:1000 | AF1062 | RRID:AB_2236897 |
| Rabbit anti-PDGFRα | Dr. William Stallcup* | 1:1000 | | |
| Rabbit anti-Phospho-Pdgfra | Cell Signaling Technology | 1:1000 | 3170 | RRID:AB_2162348 |
| **Secondary antibodies** |  |  |  |  |
| Alexa 488-donkey anti-chicken IgY | Jackson ImmunoResearch | 1:1000 | 703-545-155 | RRID:AB_2340375 |
| Cy3-donkey anti-mouse IgG (H+L) | Jackson ImmunoResearch | 1:500 | 715-165-150 | RRID:AB_2340813 |
| Cy3-goat anti-mouse IgG 2a | Jackson ImmunoResearch | 1:500 | 115-165-206 | RRID:AB_2338695 |
| Alexa 647-goat anti-mouse IgG 2a | Jackson ImmunoResearch | 1:200 | 115-605-206 | RRID:AB_2338917 |
| Cy3-goat anti-mouse IgG 2b | Jackson ImmunoResearch | 1:500 | 115-165-207 | RRID:AB_2338696 |
| Alexa 647-goat anti-Mouse IgG 2b | Jackson ImmunoResearch | 1:200 | 115-605-207 | RRID:AB_2338918 |
| Alexa 488-donkey anti-rat IgG (H+L) | Jackson ImmunoResearch | 1:1000 | 712-545-150 | RRID:AB_2340683 |
| Alexa 488-chicken anti-rat IgG (H+L) | Thermo Fisher Scientific | 1:1000 | A-21470 | RRID:AB_2535873 |
| Cy3-bovine anti-goat IgG (H+L) | Jackson ImmunoResearch | 1:500 | 805-165-180 | RRID:AB_2340880 |
| Cy3-donkey anti-goat IgG (H+L) | Jackson ImmunoResearch | 1:500 | 705-165-147 | RRID:AB_2307351 |
| Alexa 647-donkey anti-goat IgG (H+L) | Jackson ImmunoResearch | 1:200 | 705-165-147 | RRID:AB_2307351 |
| Brilliant violet 421-donkey anti-rabbit IgG (H+L) | Jackson ImmunoResearch | 1:200 | 711-675-152 | RRID:AB_2651108 |
| Cy3-donkey anti-rabbit IgG (H+L) | Jackson ImmunoResearch | 1:500 | 711-165-152 | RRID:AB_2307443 |
| IRDye 800CW donkey anti-rabbit IgG | LI-COR | 1:15,000 | 926-32213 | RRID:AB_621848 |
| IRDye 680RD donkey anti-goat IgG | LI-COR | 1:5000 | 926-68074 | RRID:AB_10956736 |

**Table 1. Primary and secondary antibodies used**
| **Antibody** | **Source** | **dilution** | **cat no** | **RRID** |
| --- | --- | --- | --- | --- |
| **Primary antibodies** |  |  |  |  |
| Mouse anti-CC1 | Millipore | 1:200 | OP80 | RRID:AB_2057371 |
| Rat anti-CD140a, clone APA5 | BD Sciences |  | 558774 | RRID:AB_397117 |
| Rat anti-CD68 | Biolegend | 1:1000 | 137001 | RRID:AB_2044003 |
| Rat anti-F4/80 | Bio-Rad | 1:200 | MCA497R | RRID:AB_323279 |
| Chicken anti-GFP | Aves Laboratories | 1:500 | GFP-1010 | RRID:AB_2307313 |
| Mouse anti-glyceraldehyde-3-phosphodehydrogenase (GAPDH) | Millipore | 1:2000 | MAB374 | RRID:AB_2107445 |
| Goat anti-Iba1 | Invitrogen | 1:500 | PA5-18039 | RRID:AB_10982846 |
| Rabbit anti-laminin | Sigma | 1:1000 | L8271 | RRID:AB_477162 |
| Mouse anti-myelin basic protein (MBP, SMI99) | Biolegend | 1:2000 | 808403 | RRID:AB_2734562 |
| Mouse anti-Neurofilament H (nonphosphorylated, SMI32; 1:3000) | Biolegend | 1:3000 | 801701 | RRID:AB_2564642 |
| Rabbit anti-NG2 | Millipore | 1:500 | AB5320 | RRID:AB_11213678 |
| Goat anti-Nrp1 | R&D Systems | 1:200 | AF566 | RRID:AB_355445 |
| Mouse anti-Olig2 | Millipore | 1:500 | MABN50 | RRID:AB_10807410 |
| Goat anti-PDGFRα | R&D Systems | 1:1000 | AF1062 | RRID:AB_2236897 |
| Rabbit anti-PDGFRα | Dr. William Stallcup* | 1:1000 | | |
| Rabbit anti-Phospho-Pdgfra | Cell Signaling Technology | 1:1000 | 3170 | RRID:AB_2162348 |
| **Secondary antibodies** |  |  |  |  |
| Alexa 488-donkey anti-chicken IgY | Jackson ImmunoResearch | 1:1000 | 703-545-155 | RRID:AB_2340375 |
| Cy3-donkey anti-mouse IgG (H+L) | Jackson ImmunoResearch | 1:500 | 715-165-150 | RRID:AB_2340813 |
| Cy3-goat anti-mouse IgG 2a | Jackson ImmunoResearch | 1:500 | 115-165-206 | RRID:AB_2338695 |
| Alexa 647-goat anti-mouse IgG 2a | Jackson ImmunoResearch | 1:200 | 115-605-206 | RRID:AB_2338917 |
| Cy3-goat anti-mouse IgG 2b | Jackson ImmunoResearch | 1:500 | 115-165-207 | RRID:AB_2338696 |
| Alexa 647-goat anti-Mouse IgG 2b | Jackson ImmunoResearch | 1:200 | 115-605-207 | RRID:AB_2338918 |
| Alexa 488-donkey anti-rat IgG (H+L) | Jackson ImmunoResearch | 1:1000 | 712-545-150 | RRID:AB_2340683 |
| Alexa 488-chicken anti-rat IgG (H+L) | Thermo Fisher Scientific | 1:1000 | A-21470 | RRID:AB_2535873 |
| Cy3-bovine anti-goat IgG (H+L) | Jackson ImmunoResearch | 1:500 | 805-165-180 | RRID:AB_2340880 |
| Cy3-donkey anti-goat IgG (H+L) | Jackson ImmunoResearch | 1:500 | 705-165-147 | RRID:AB_2307351 |
| Alexa 647-donkey anti-goat IgG (H+L) | Jackson ImmunoResearch | 1:200 | 705-165-147 | RRID:AB_2307351 |
| Brilliant violet 421-donkey anti-rabbit IgG (H+L) | Jackson ImmunoResearch | 1:200 | 711-675-152 | RRID:AB_2651108 |
| Cy3-donkey anti-rabbit IgG (H+L) | Jackson ImmunoResearch | 1:500 | 711-165-152 | RRID:AB_2307443 |
| IRDye 800CW donkey anti-rabbit IgG | LI-COR | 1:15,000 | 926-32213 | RRID:AB_621848 |
| IRDye 680RD donkey anti-goat IgG | LI-COR | 1:5000 | 926-68074 | RRID:AB_10956736 |

### Transmission Electron Microscopy

After perfusion and post-fixation of LPC-injected mice at day 28 dpl, brains were washed in PBS. 100-μm-thick coronal slices were obtained by slicing the brains with a Leica vibratome VT1000S. Slices containing the LPC-induced lesion were selected and fixed again in 4% PFA and 2% glutaraldehyde in 0.1M sodium phosphate buffer for 2 hours. The corpus callosum within the region of interest was dissected and processed for resin embedding. Specimens were further fixed with 1% Osmium tetroxide at room temperature for 2h. Tissue was dehydrated through a series of graded ethanol, during which 1.5% uranyl acetate was included. 100% ethanol was gradually mixed with an increasing portion of propylene oxide and finally incubated in propylene oxide. Subsequently, infiltration with Spurr’s resin was carried out with an increasing portion of resin diluted with propylene oxide. Infiltration was completed with 100% epon resin. Tissue was embedded in flat double end molds and polymerized at 70°C for 48 hours. Semithin sections were cut with a Diatome™ diamond knife on a Leica Ultracut UCT microtome and stained with methylene blue/azure blue. The region of interest was selected with upright Leica DMR and Zeiss AxiovertM200 inverted microscopes, and ultrathin sections were cut with an ultra 45° Diatome TM diamond knife. Sections were collected on 100 mesh copper grids. Images were obtained using a bright field FEI Tecnai Biotwin G2 Spirit (Netherlands) transmission electron microscope operated at an accelerating voltage of 80 kV and equipped with an AMT 2k (4 megapixel) XR40 CCD camera.

### Slice cultures

Slice cultures from the forebrain and cerebellum were prepared from P8 NG2cre;Z/EG mice as previously described ^15,29^ or from P8 mg-Nrp1-cont or cko mice after 4OHT injection at P2-3 in vivo. 300-μm thick slices were maintained at air-liquid interface on Millicell cultures inserts (Millipore) and incubated in slice media for 7 days, with media change every other day. On day 7, PDGF AA (R&D Systems, 221-AA) was added to the cultures and the slices were incubated for 48 hours. During the last 5 hours of incubation, 10 μM EdU was added to the slice medium. To block Nrp1 or PDGFRα function, slices were incubated in different concentrations of goat anti-mouse Nrp1 antibody (R&D Systems, AF566) or 1 μg/mL of goat anti-mouse PDGFRα antibody (R&D Systems, AF1062) during the last 48 hours of incubation (on day 7). Nrp1-Fc fusion protein containing the extracellular domain of rat Nrp1 (amino acid 22-854 minus 811-828; R&D Systems, 566-NNS) or the control human Fc dimer (R&D Systems, 100-HG) was added to slice cultures at the indicated concentrations for the last 48 hours of incubation. At the end of the incubation, slices were fixed with 4% paraformaldehyde and processed for immunofluorescence labeling and EdU detection.

### Dissociated OPC cultures

OPCs from the cerebral cortex of P4-5 wild type CD1 mice were immunopanned for PDGFRα as previously described ^30^, using CD140a rat anti-mouse PDGFRα antibody. Purified OPCs were plated on glass coverslips coated with poly-D-lysine (Sigma, P7405, 10 μg/mL) at a density of 40,000 cells/well in 24-well plates for EdU incorporation assays or on 35-mm tissue culture dishes coated with poly-L-lysine (Sigma, P1524; 30 μg/mL) at a density 10×10^6^ cells/dish for immunoblotting. For EdU detection, the cells on coverslips were initially incubated in DMEM-Sato medium and PDGF AA for 2 days, and 10 μM EdU was added to the cells for 6 hours prior to staining for Olig2, NG2, and EdU. For detecting PDGFRα phosphorylation by immunoblotting, immunopurified OPCs were incubated with DMEM-Sato medium without PDGFAA for 12 h.Then, the medium was replaced with new medium containing 2 μg/mL Nrp1-Fc or control Fc protein and 15 ng/mL PDGFAA and incubated at 37°C for 30 min, after which the cells were promptly chilled on ice for protein extraction.

For Nrp1-Fc-PDGFRα capping experiments, immunopanned OPCs from P4-5 CD1 mice were plated on PDL coated coverslips at a density of 40,000-50,000 cells / well in 24-well plates and incubated in DMEM-Sato medium overnight. Then, 2 μg/mL control human IgG1-Fc (cont-Fc) or Nrp1-Fc was added in the presence of 15ng/mL PDGF AA and incubated at 4°C or 36°C for 30min. The coverslips were washed with cold PBS, fixed with 4%PFA, and then immunolabeled with mouse anti-Olig2 (Millipore MABN50, 1:500) and rabbit anti-PDGFRα antibodies (obtained from Dr William Stallcup, Sanford Burnham Institute, La Jolla, CA; 1:1,000 dilution). Following washes, cells were incubated in Alexa488-goat anti-human Fc, Cy3-bovine anti-rabbit, and Alexa647-donkey anti-mouse antibodies.

### Immunoblotting

Purified immunopanned OPCs were rinsed with chilled PBS containing 1x PhosSTOP (Sigma-Aldrich-Roche, 4906845001) and 1x halt protease inhibitor cocktail (ThermoFisher, 87786) and harvested in 1 mL of the same buffer. The cell pellet was lysed by homogenizing in RIPA Buffer (ThermoFisher) supplemented with PhosSTOP and protease inhibitors on ice for 30 minutes and then centrifuged to remove insoluble matter. The cleared lysate was frozen in liquid nitrogen and stored at −80°C. Protein concentration was measured using DC protein assay (Bio-Rad). Samples were denatured with 4X Bolt LDS sample buffer and 10X sample reducing agent (ThermoFisher B0007 and B0004, respectively) and heated at 70°C for 10 minutes. Samples (30 μg/lane) and prestained molecular weight standards (LI-COR, 928-7000) were electrophoresed through 8% Bolt Bis-Tris plus polyacrylamide gels (ThermoFisher) in Bolt MOPS SDS running buffer (ThermoFisher). Proteins were transferred to Immobilon FL membranes (EMD-Millipore) using a transfer buffer containing 1.25 mM bicine, 1.25 mM bis-Tris, and 0.05 mM EDTA. Blots were blocked in Odyssey TBS (Tris-buffered saline) blocking buffer (LI-COR) at room-temperature for 1h. Blots were incubated at 4°C overnight in primary antibodies diluted in TBS blocking buffer containing 0.1% Tween 20. The primary antibodies were rabbit anti-Phospho-PDGFRα (Cell signaling) and goat anti-mouse PDGFRα that recognizes both phosphorylated and non-phosphorylated forms (R&D Systems). Blots were washed with TBST buffer containing 137 mM NaCl, 20 mM Tris HCl, pH 7.6, and 0.1% Tween 20) 4 times, 10 minutes each, and incubated with IRDye 800RD donkey anti-rabbit or IRDye 680RD donkey anti-goat secondary antibodies diluted in Odyssey blocking buffer with 0.1%Tween 20 and 0.01%SDS. Blots were washed with TBST and imaged on a LI-COR Odyssey Imager. To quantify protein expression, the density of protein bands was determined using Image Studio Lite (LI-COR).

### Fluorescence microscopy and quantification

Images of fluorescently labeled tissue sections, slice cultures, and dissociated cells were acquired using Leica SP8 confocal microscope (Advanced Light Microscopy Facility) or Zeiss Axiovert 200M with apotome. Images were analyzed with the Leica LAS X, Zeiss Axiovision, or ImageJ. For quantification, images were captured from several random fields of defined area within the cortex, corpus callosum, or the cerebellar white matter, based on DAPI label and blind to the experimental labels, and immunolabeled cells in each field were counted.

To quantify the proliferative OPCs that were in contact with microglia, 0.35-μm *z*-stack images were acquired from 20-μm-thick sections using a 40X objective on a Leica SP8 confocal microscope at 1024 × 1024 pixel image size and 700 Hz scan speed. Three-dimensional images were reconstructed from the *z*-stacks using the 3D module of the Leica LAS X software. Each PDGFRα+ OPC was marked and first examined for contact with Nrp1+ microglia by rotating the plane in all directions. Then, the EdU channel was revealed, and the OPC was scored for whether it had EdU label.

For quantification of the degree of myelination, sections from P14 corpus callosum or demyelinated corpus callosum at 28 dpl were immunolabeled for MBP, and MBP immunofluorescence intensity was quantified on images scanned on the Leica SP8 confocal microscope. Fiji (ImageJ, version 1.53a) was used to obtain the average integrated pixel density of gray values over a defined region of interest. Background values were obtained from areas devoid of signal.

For dissociated OPCs, images were taken from randomly selected fields from each coverslip based on DAPI label and the labeled cells were quantified. Adobe Photoshop was used to generate image panels for the figures. Quantification data are represented as means ± standard deviations. The specific statistical method used to evaluate each dataset is indicated in the figure legends.

## RESULTS

### Nrp1 modulates PDGF AA-dependent OPC proliferation in white matter

To determine whether the proliferative response of OPCs to PDGF AA was modulated by Nrp1, we tested the effects of anti-Nrp1 antibody on PDGF AA-induced OPC proliferation in forebrain slice cultures from postnatal day 8 (P8) NG2cre;Z/EG double transgenic mice ^15^ (**Fig 1A**). In the presence of 50 ng/mL PDGF AA alone, 42% of EGFP+ OPCs in the corpus callosum were EdU+ (**Fig 1D, F**). By contrast only 18.4% of those in the cortex were EdU+ in the presence of PDGF AA (**Fig 1B, F**). Anti-Nrp1 antibodies elicited a dose-dependent decrease in the proportion of EGFP+ cells that proliferated in response to PDGF AA in the corpus callosum (**Fig 1F**), but there was no effect on basal OPC proliferation in gray matter (**Fig 1C, F**). We did not observe any increase in TUNEL+ cells in the anti-Nrp1-treated slices (**Fig 1G, H**) compared to slices treated with a control IgG, indicating that the antibody did not cause significant cell death. Thus, Nrp1 appeared to be necessary for PDGF AA-mediated OPC proliferation in the corpus callosum.

**Fig 1.**
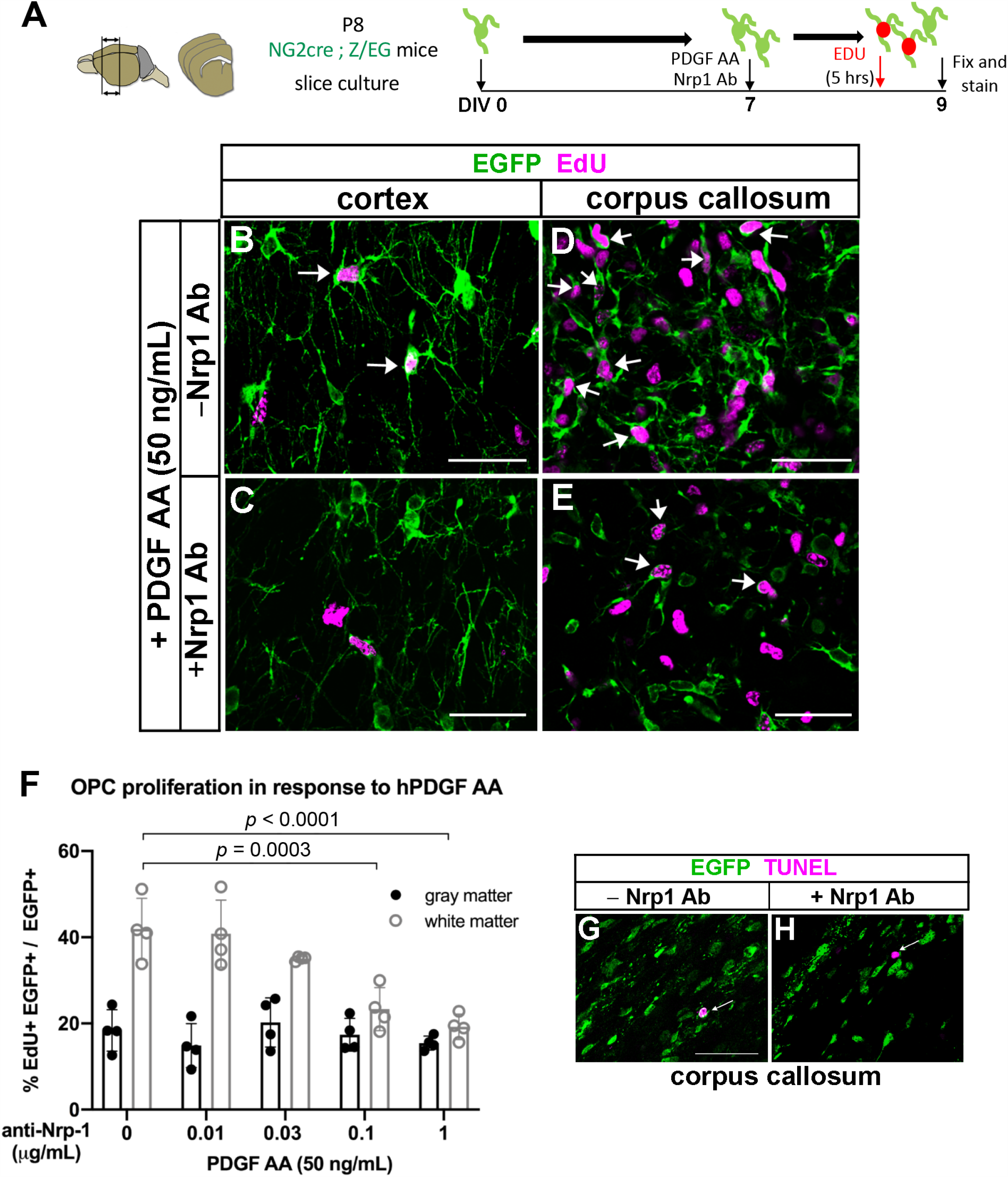
Anti-Nrp1 antibody blocks PDGF AA-mediated OPC proliferation in white but not gray matter. **A**. Schematic showing slice cultures from P8 NG2cre;Z/EG mice to assay for OPC proliferation in response to 50 ng/mL PDGF AA and different concentrations of anti-Nrp1 antibody. **B-E**. Slice cultures from P8 NG2cre;Z/EG mice that were fixed and labeled for EGFP and EdU in the cortex (B and C) or corpus callosum (D and E) in the presence of 50 ng/mL PDGF AA in the absence (B and D) or presence (C and E) of 1μg/mL anti-Nrp1 antibody. Arrows, examples of EGFP+ EdU+ cells. Scale bars, 50 μm. **F**. Quantification of OPC proliferation in slice cultures after 5 hours of EdU labeling. *y*-axis, proportion of EGFP+ cells that were EdU+. Tukey’s multiple comparisons test, *n*=4, F(4, 30) = 9.966, means ± standard deviations. The differences between OPC proliferation in gray and white matter were significant at anti-Nrp1 antibody concentrations of 0 (*p* < 0.0001) and 0.01 μg/mL (*p* = 0.0065), but not at higher concentrations of the antibody. Two-way ANOVA, Tukey’s multiple comparisons test, F(1, 30) = 89.71, *n*=4. **G-H**. Slice cultures treated with 1 μg/mL control goat IgG (G) or goat anti-Nrp1 antibody (H) and labeled for EGFP and TUNEL (magenta). Arrows indicate a TUNEL+ cell. Scale bars, 50 μm.

### Nrp1 is expressed on amoeboid microglia in the developing corpus callosum

To determine the relevant source of Nrp1 that could modulate PDGF AA-dependent OPC proliferation in the corpus callosum, we examined the developing mouse CNS tissues for Nrp1 expression. In P5 brain, Nrp1 was detected widely in the forebrain (**Fig 2A**), including vascular cells (arrowheads in Fig 2A, C) along laminin+ blood vessels (**Fig S1A**), as expected from previous reports on Nrp1 on vascular endothelial cells ^22^. Nrp1 immunoreactivity was also detected on axons in the dorsal spinal cord (**Fig S1B**) in P5 mice and in the dorsal corpus callosum in E18.5 brain (**Fig S1C**), as previously shown (Kawakami et al. 1996). In addition, there was strong cellular staining in a subregion of the corpus callosum (**Fig 2A**, boxed area magnified in **Fig 2D-F**). The majority of the round Nrp1+ cells in the corpus callosum also expressed the microglial cell surface antigen F4/80 ^31^ and had the appearance of amoeboid microglia (**Fig 2A-F**). Similarly, we detected Nrp1 on a subpopulation of F4/80+ amoeboid microglia in P5 cerebellar white matter (not shown). We did not detect Nrp1 on PDGFRα+ OPCs (**Fig 2G-I**), although Nrp1+ microglial processes were closely apposed to OPC cell bodies and processes (arrows in **Fig 2I**). The close proximity to Nrp1-expressing amoeboid microglia to OPCs suggested that microglial Nrp1 in the white matter could be modulating the proliferative response of OPCs to PDGF AA.

**Fig 2.**
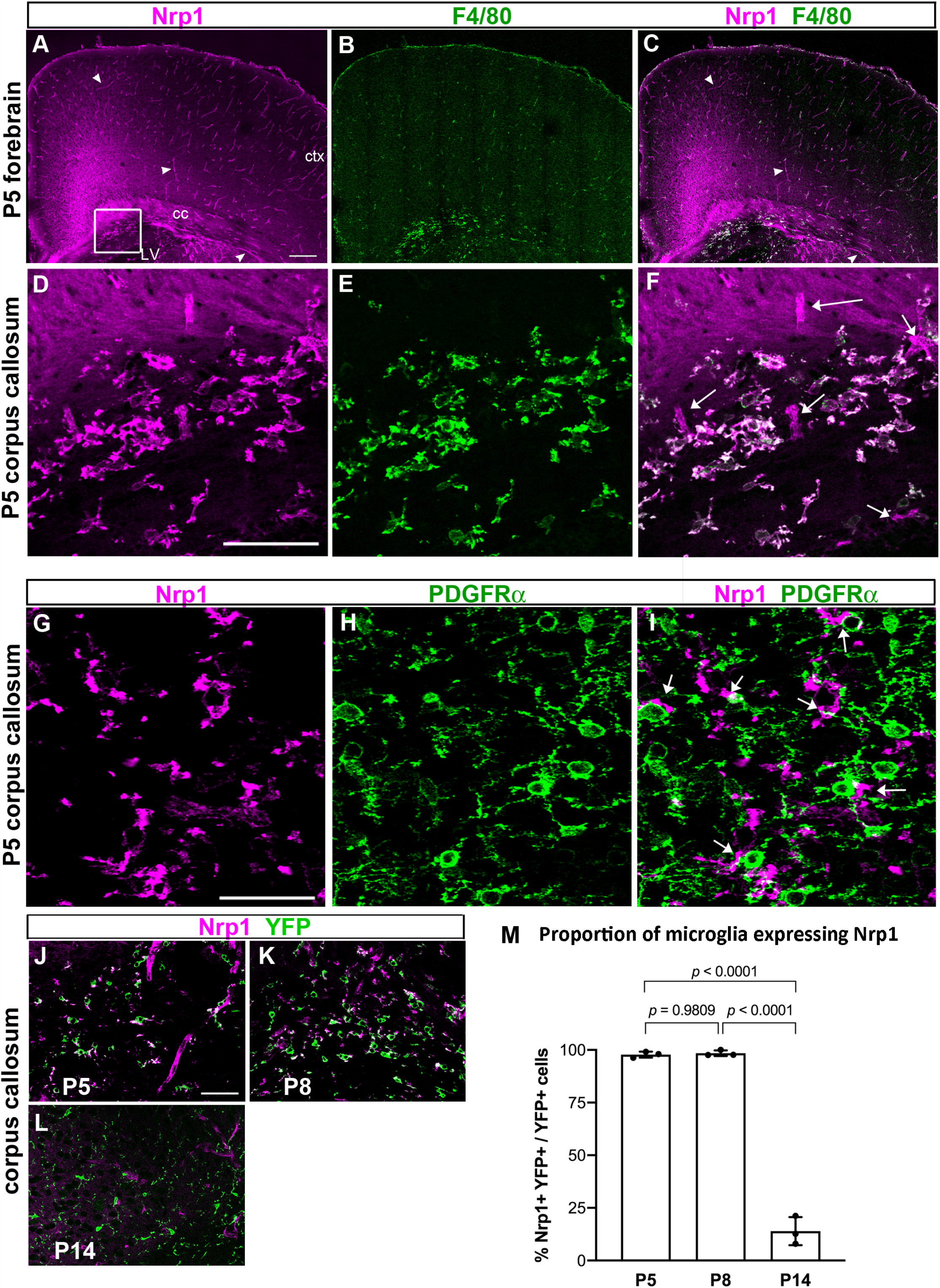
Nrp1 expression in the developing postnatal brain. **A-C**. Low magnification view of a coronal forebrain section labeled for Nrp1 and F4/80. Arrowheads, examples of Nrp1+ blood vessels. Arrowheads, blood vessels. Boxed area with a cluster of Nrp1+ cells is shown in D-F. scale, 100 μm. ctx, cortex; cc, corpus callosum; LV lateral ventricle. **D-F**. High magnification view showing Nrp1+ cells in the corpus callosum express F4/80. Arrows in F indicate blood vessels that are F4/80-negative. Scale, 50 μm. **G-I**. Double labeling for Nrp1 and PDGFRα in P5 corpus callosum. Nrp1 is not expressed on PDGFRα+ OPCs, but Nrp1+ processes are found close to OPC cell bodies and processes (arrows in I). **J-L**. Double labeling for Nrp1 and YFP in Cx3CR1^creERT2-ires-EYFP^ mice from P5 through P14. Scale = 50 μm. **M**. The proportion of EYFP+ microglia that expressed Nrp1 in the corpus callosum. One-way ANOVA, Tukey’s multiple comparisons test, *n* = 3, F (2, 6) = 438.8.

To further examine the enrichment of Nrp1 among microglia in white matter, we examined the expression of Nrp1 on EYFP+ cells in the neocortex and corpus callosum of Cx3Cr1^creERT2-ires-EYFP^ mice from P5 to P30 (**Fig 2J-M, Fig S2**). At P5 and P8, Nrp1 was expressed on the majority of EYFP+ amoeboid microglia in the corpus callosum (97.8% at P5 and 98.4% at P8), but the enrichment declined by P14 to 14.0% (**Fig 2M**), and Nrp1 was no longer detected at P30, as the EYFP+ microglia in the corpus callosum displayed progressively more ramified morphology. In the cortex, EYFP+ microglia had ramified morphology from P5 through P30 and did not co-express Nrp1. Thus, Nrp1 was preferentially expressed on amoeboid cells in the early postnatal white matter.

The amoeboid microglia that were found in P5 corpus callosum also expressed CD68 (**Fig S3, A-D**, arrows), which is a lysosomal marker of phagocytosis ^32,33^ and hence an indicator of phagocytic activity. The microglia in P14 corpus callosum exhibited a very small amount of punctate immunoreactivity for CD68, which was significantly lower compared to that in P5 corpus callosum, and these microglia were Nrp1-negative (**Fig S3, E-H**, arrowheads)

### Microglia-specific Nrp1 deletion reduces developmental OPC proliferation in white matter

To examine the effects of microglial Nrp1 on OPC proliferation, we deleted Nrp1 specifically from microglia using Nrp1 conditional knockout mice ^26^ crossed to Cx3CR1^creERT2-ires-EYFP^ mice, which express tamoxifen-inducible cre specifically in Cx3CR1-expressing microglia ^27^ (**Fig 3A**). When we induced Nrp1 deletion in microglia in mg-Nrp1-cont (fl/+) and mg-Nrp1-cko (fl/fl) mice by injecting 4-hydroxytamoxifen (4OHT) at P2 and P3 and analyzed at P5, none of the EYFP+ microglia in mg-Nrp1 cko mice had detectable Nrp1 expression in the corpus callosum, whereas Nrp1 was detected on >99% of EYFP+ microglia in mg-Nrp1 cont heterozygous mice (**Fig S4 A-D**). Microglial Nrp1 deletion did not affect the density of microglia or astrocytes (**Fig S4 E-F**). Nrp1 immunoreactivity on blood vessels remained detectable in mg-Nrp1-cko (**Fig S4C**, arrow), and the relative area in the corpus callosum occupied by blood vessels was not altered in mg-Nrp1-cko (**Fig S4G**).

**Fig 3.**
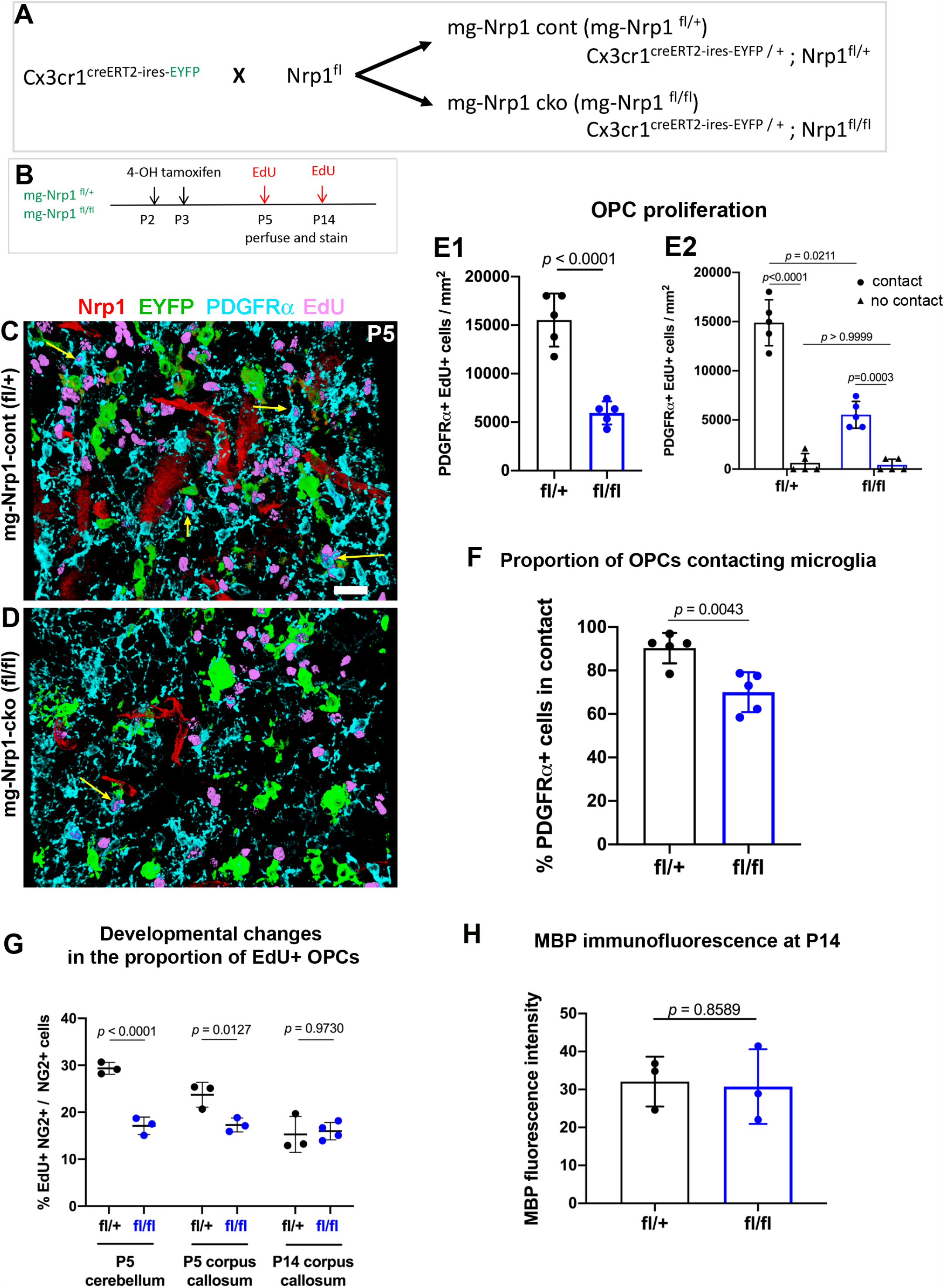
Effects of mg-Nrp1 cko on OPC proliferation in the white matter. **A**. Schematic of generating microglia-specific Nrp1 knock out (mg-Nrp1-cko) and the heterozygous control (mg-Nrp1-cont). **B**. Schematic of tissue analysis at P5 and P14 after cre induction at P2-3. **C-D**. 3D images of P5 corpus callosum labeled for Nrp1, EYFP, PDGFRα, and EdU. Scale, 20μm. Arrows, examples of EdU+ PDGFRα+ cells in contact with EYFP+ microglia. **E**. Quantification of proliferation of OPCs in mg-Nrp1-cont and cko. **E1**. Total EdU+ OPC density in P5 corpus callosum. Unpaired Student’s t-test, *n* = 5, t = 7.171, df = 8. **E2**. The density of EdU+ OPCs in cont or cko with (circles) or without (triangles) contact with microglia. Two-way ANOVA, Sidak’s multiple comparisons test,F(1, 16) = 53.41, *p* < 0.0001, *n* = 5. **F**. Proportion of PDGFRα+ OPCs that were in contact with EYFP+ microglia. Unpaired Student’s t-test, *n* = 5, t = 3.940, df = 8. **G**. Proportion of OPCs that were EdU+ in P5 cerebellum, P5 corpus callosum, and P14 corpus callosum. Two-way ANOVA, Sidak’s multiple comparisons test. F(1, 13) = 32.44, *p* < 0.0001, *n*= 3-4. **H**. Quantification of MBP immunofluorescence intensity in P14 corpus callosum of cont and cko mice. N = 3, t = 0.1896, df = 4.

Having confirmed microglia-specific deletion of Nrp1 with this model, we next examined OPC proliferation in the corpus callosum of mg-Nrp1-cont and cko mice by EdU pulse labeling at P5 after microglial Nrp1 deletion at P2 and P3 (**Fig 3B**). Comparison of the density of PDGFRα+ EdU+ cells in mg-Nrp1-cont and cko corpus callosum revealed a 2.6-fold reduction in cko (**Fig 3 C-D, E1**). Since we observed that OPCs were closely apposed to Nrp1+ microglia in P5 corpus callosum as described above (Fig 2 G-I), we investigated whether OPCs that were in contact with Nrp1+ microglia were proliferating more than OPCs without microglial contact. We used 3D rotation of confocal *z*-stacks from P5 sections labeled for PDGFRα, Nrp1, and EYFP, and EdU and assessed whether each PDGFRα+ OPC had an EYFP+ microglial element directly apposed to it and whether it had incorporated EdU (**Fig 3 C-D**, arrows). Among the proliferating OPCs, 95.9% in mg-Nrp1-cont and 92.8% in mg-Nrp1-cko had contact with microglia. There was no significant difference in the density of proliferating OPCs that were not in contact with microglia between the two genotypes (**Fig 3E2**, triangles). By contrast, the density of proliferating OPCs that were in contact with microglia was 2.7-fold lower in mg-Nrp1-cko compared to cont (**Fig 3E**, circles). This indicated that the decrease in OPC proliferation in the corpus callosum of P5 mg-Nrp1-cko was largely due to the decrease in OPCs that were in contact with microglia. In mg-Nrp1-cko, the proportion of total PDGFRα+ OPCs, regardless of EdU incorporation, that were contacting microglia was 22% lower than that in mg-Nrp1-cont (**Fig 3F**), suggesting that the presence of Nrp1 on microglia could be facilitating OPC contact. We also observed a similar reduction in the proliferation of OPCs contacting microglia in P5 cerebellum (data not shown).

We next examined whether the decrease in OPC proliferation in the white matter of mg-Nrp1-cko mice was restricted to the period in which Nrp1 was highly expressed on amoeboid microglia. While the proportion of NG2+ OPCs that had incorporated EdU+ was significantly lower in the cerebellar white matter and corpus callosum of P5 mg-Nrp1-cko, the difference was no longer seen in P14 corpus callosum (**Fig 3G**). Thus, the developmental age during which microglial Nrp1 deletion reduced OPC proliferation coincided with the temporal window during which Nrp1 was expressed on amoeboid microglia in the corpus callosum (**Fig 2M, Fig S2**).

The density of total OPCs in the corpus callosum of mg-Nrp1-cko mice was 1.7-fold lower than that in control mice at P5 and remained 1.4-fold lower at P14. TUNEL labeling did not reveal any TUNEL+ cells in the corpus callosum of mg-Nrp1-cont or mg-Nrp1-cko at P5 (**Fig S4H-J**), suggesting that microglial deletion of Nrp1 did not increase cell death. The transient reduction in OPC density did not lead to long-lasting changes in myelination, as judged by comparable levels of myelin basic protein (MBP) immunofluorescence in the corpus callosum in P14 mg-Nrp1-cont and cko mice (**Fig 3H, Fig S4 K-P**).

To further examine whether microglial deletion of Nrp1 compromised the proliferation of OPCs to PDGF AA, we prepared slice cultures from P8 mg-Nrp1-cont and mg-Nrp1-cko mouse forebrains after 4OHT administration at P2-3 and examined the proliferation of OPCs in gray and white matter in response to 50 ng/mL PDGF AA (**Fig S5**). There was no effect of Nrp1 deletion on EdU incorporation into gray matter OPCs. By contrast, EdU incorporation into OPCs in the corpus callosum of mg-Nrp1-cko slices was reduced to 57% of mg-Nrp1-cont.

### Microglial Nrp1 deletion compromises OPC expansion after demyelination

Our immunolocalization studies revealed that Nrp1 was abundantly expressed on amoeboid microglia that appeared transiently in the corpus callosum during the first postnatal week and was downregulated on microglia by P14 as they became more ramified. To determine whether Nrp1 expression could be upregulated in pathological conditions that are known to increase OPC proliferation, we used an acute chemically induced demyelination model created by injecting α-lysophosphatidylcholine (LPC, lysolecithin) into the corpus callosum of 2-to 3-month-old mg-Nrp1-cont and mg-Nrp1-cko mice (**Fig 4A**). Nrp1 was deleted in microglia by tamoxifen injection prior to induction of demyelination, with a deletion efficiency of 99.5% of YFP+ cells.We first examined whether Nrp1 was re-expressed on microglia after demyelination in mg-Nrp1-cont mice. Three days after PBS injection, EYFP+ cells surrounding the injection site had the morphology of resting ramified microglia and did not express Nrp1 (**Fig 4B**). By contrast, three days after LPC injection (3 dpl), there was strong activation of microglia, and Nrp1 was robustly upregulated on the activated EYFP+ microglia/macrophages (**Fig 4C**). At 7 dpl, EYFP+ activated microglia/macrophages were abundantly detected and expressed the lysosomal protein CD68 in the demyelinated lesion of both mg-Nrp1-cont and mg-Nrp1-cko corpus callosum (**Fig 4D-E**). In mg-Nrp1-cont lesions, Nrp1 immunoreactivity was detected on the surface of CD68+ macrophages (**Fig 4D**, arrows). This indicated that Nrp1 was highly upregulated on activated microglia/macrophages that appeared in the demyelinated corpus callosum. In the cko lesions, the majority of the CD68+ cells lacked Nrp1 (**Fig 4E**, arrowheads). While Nrp1 was undetectable in 99.5% of YFP+ cells at 3 dpl, we detected a small number of CD68+ YFP-cells that expressed Nrp1 in the cko corpus callosum at 7 dpl (**Fig 4E**, arrows), which were likely to have entered the lesion from a Cx3cr1-negative precursor. The density of EYFP+ microglia/macrophages was similar in mg-Nrp1-cont and mg-Nrp1-cko mice at 3 dpl **(Fig 4F)**, indicating that the lack of Nrp1 on microglia in mg-Nrp1-cko lesions did not affect their infiltration or activation.

**Fig 4.**
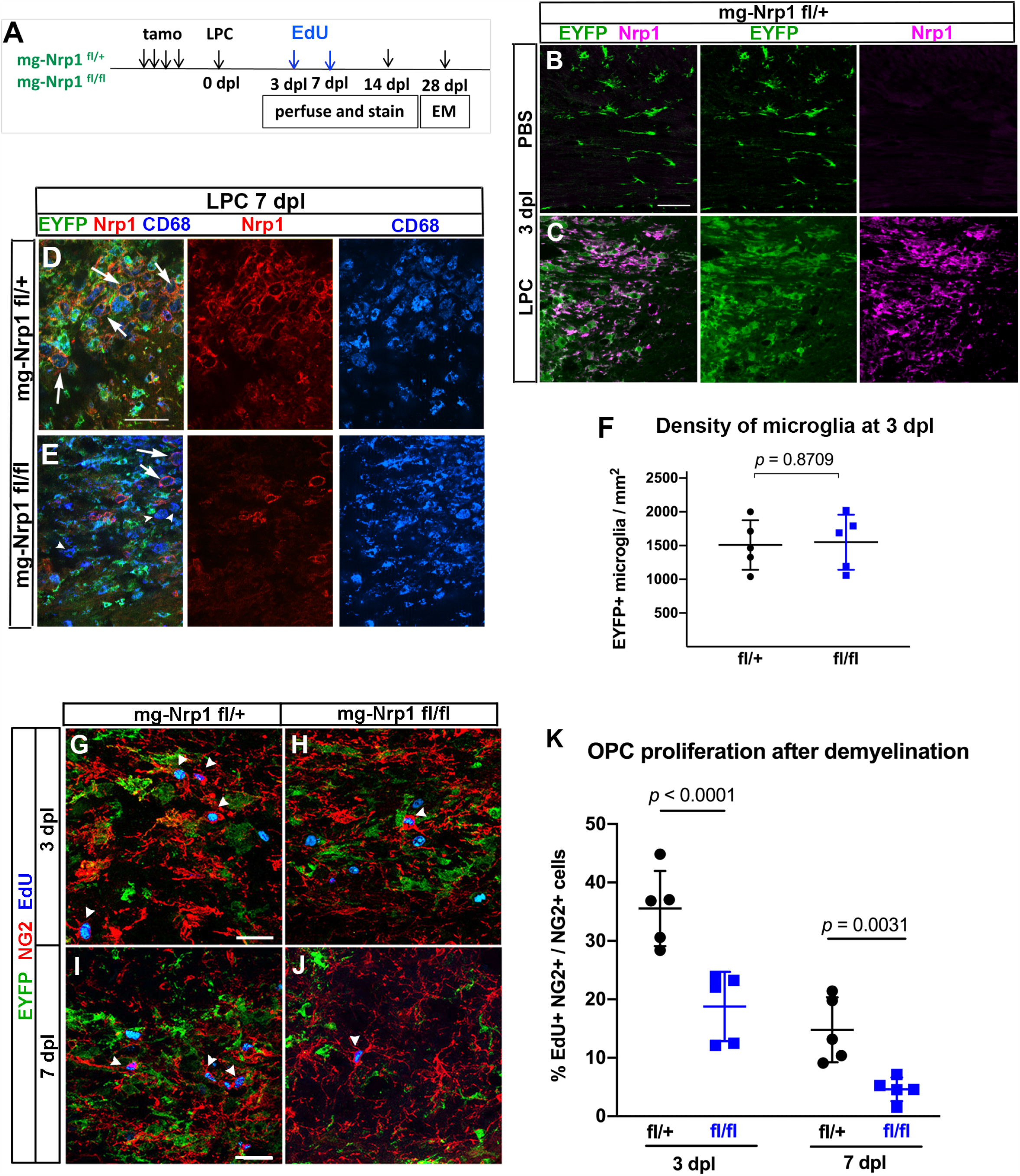
Effects of mg-Nrp1-cko on OPC proliferation and myelin repair after demyelination. **A**. Schematic of demyelination experiments. **B-C**. Nrp1 upregulation on EYFP+ microglia/macrophages 3 days after LPC injection but not after PBS injection. Scale, 50 μm. **D-E**. Similar extent of CD68 activation and infiltration of activated microglia/macrophages into demyelinated lesion at 7 dpl in mg-Nrp1-cont (**D**) and mg-Nrp1-cko (**E**) mice. Arrows, Nrp1+ CD68 macrophages; arrowheads, Nrp1-negative CD68+ cells. Scale, 50 μm. **F**. Quantification of the density of EYFP+ microglia in the lesion at 3 dpl. Unpaired Student’s t-test, t = 0.1678, df = 8, *n* = 5. **G-J**. Labeling for EdU, NG2, and EYFP in mg-Nrp1-cont and cko mice sacrificed at 3 dpl after EdU pulse labeling. Arrowheads, NG2+ EdU+ OPCs. Scale, 25 μm. **K**. Quantification of proliferating OPCs at 3 and 7 dpl in cont (black) and cko (blue) mice showing significantly lower extent of EdU incorporation into OPCs in mg-Nrp1-cko mice. OPC proliferation was higher at 3 dpl compared to 7 dpl for both genotypes (*p* = 0.0007 for cont and *p*= 0.0351 for cko). Two-way ANOVA, Tukey’s multiple comparisons test, F(1, 16) = 54.70 for comparison between fl/+ and fl/fl, F(1, 16) = 32.52 for comparison between 3 and 7 dpl, *n*=5.

At 3 and 7 dpl, we pulse-labeled LPC-injected mice with EdU and examined the effect of microglial Nrp1 deletion on OPC proliferation in the lesion. EdU incorporation into EYFP-negative, NG2+ OPCs at the lesion site in mg-Nrp1-cko was 1.9-fold lower than that in mg-Nrp1-cont lesions at 3 dpl and 3.2-fold lower at 7 dpl (**Fig 4F-K**). OPC proliferation was 2-to 3-fold higher at 3 dpl compared to 7 dpl for both genotypes. These observations indicate that loss of Nrp1 on activated microglia significantly compromised the ability of OPCs to proliferate in response to acute demyelination.

### Microglial Nrp1 deletion compromises OL regeneration and myelin repair

To examine whether the reduced proliferation of OPCs in mg-Nrp1-cko mice affected subsequent events in myelin regeneration, we examined the lesion at 14 dpl. Quantification of OLs in and around the lesion at 14 dpl revealed that the density of CC1+ OLs in mg-Nrp1-cko lesion was reduced to 63% of that in mg-Nrp1-cont lesion (**Fig 5 A-C**). To assess the extent of myelin repair, we first estimated the demyelinated area at 14 dpl by taking the area that exhibited reduced myelin basic protein (MBP) and elevated non-phosphorylated neurofilament immunoreactivity. The demyelinated area at 14 dpl was 1.74 times larger in the mg-Nrp1-cko mice compared with that in the mg-Nrp1-cont mice (**Fig 5 D-F**). Thus, the reduction of oligodendrocyte density in the knockout animals correlated with the larger area containing demyelinated axons, suggesting that the repair process was impaired in mg-Nrp1-cko mice.

**Fig 5.**
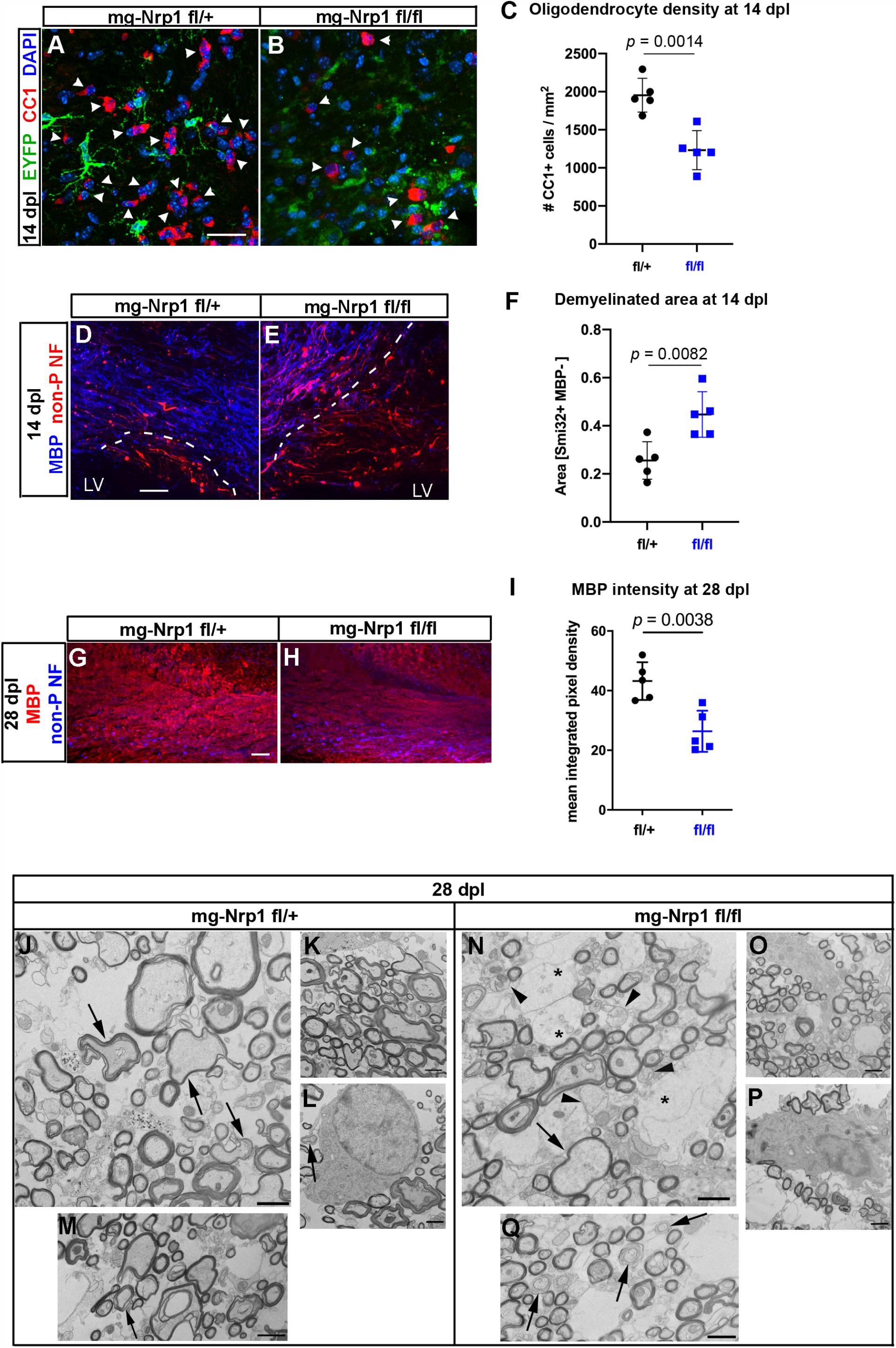
The extent of oligodendrocyte regeneration and myelin repair in mg-Nrp1-cko. **A-B**. Labeling mg-Nrp1-cont (L) and cko (M) for CC1, DAPI, and YFP at 14 dpl. Arrowheads, CC1+ cells. Scale, 25 μm. **C**. Quantification of OL density in cont and cko lesions at 14 dpl. Student’s t-test, unpaired, two-tailed, *n*=5, t = 4.753, df = 8. **D-E**. Labeling mg-Nrp1-cont (O) and cko (P) for MBP and non-phosphorylated neurofilaments (non-P-NF) using smi-32 antibody to identify axons that have not yet undergone complete remyelination. Dotted lines indicate the extent of partial or complete demyelination. Scale, 50μm. LV, lateral ventricle. **F**. Quantification of demyelinated area in cont and cko lesions at 14 dpl. Student’s t-test, unpaired, two-tailed, *n*=5, t = 3.490, df = 8. **G-H**. Labeling for MBP and non-phosphorylated neurofilaments in the lesioned corpus callosum of mg-Nrp1-cont (G) and cko (H) mice at 28 dpl.Quantification of MBP immunofluorescence in the lesioned corpus callosum of mg-Nrp1-cont and cko mice at 28 dpl. Student’s t-test, unpaired, two-tailed, *n*=5, t = 4.031, df = 8. **J-Q**. Electron microscopic images of cross-sections of LPC-lesioned corpus callosum at 28 dpl. **J-M:** mg-Nrp1-cont (fl/+) showing sheaths of various thickness (**J**, arrows), but most axons appear fully myelinated (**K**). Occasional glial processes (K) including a typical oligodendrocyte (**L**) surrounded by a few myelinated axons and one that still appears to be in the process of myelination (arrow in **L**). Occasional axons are surrounded by non-compacted myelin (**M**, arrow). **N-Q:** mg-Nr1p-cko (fl/fl) showing more axons without myelin sheaths (**N**, arrowheads), axons with thin myelin (**N**, arrow), and abundant swollen glial cell processes (**N**, *). More glial processes are seen near axons with various extent of myelination / ensheathment (**O**) and a typical oligodendrocyte surrounded by axons with varying thickness of myelin sheaths (**P**).Axons surrounded by non-compacted protrusions of glial cell processes are frequent (**O**, arrows). Scale bars 1 μm.

We next examined the impact of Nrp1 deletion from activated microglia on remyelination at 28 dpl. MBP immunofluorescence intensity in the lesioned corpus callosum of mg-Nrp1-cko mice was lower, at 61% of that in mg-Nrp1-cont mice (**Fig 5 G-I**). There was a higher level of non-phosphorylated neurofilament in the cko, which is consistent with prolonged impairment of myelin repair. To examine the extent of myelin repair in more detail, we performed ultrastructural analysis of the lesioned corpus callosum of mg-Nrp1-cont and mg-Nrp1-cko mice at 28 dpl (**Fig 5J-Q**) and searched for signs of myelin repair process. In mg-Nrp1-cont lesions, we saw that most axons were myelinated, and that some axons were surrounded by thin or less compacted myelin sheaths indicating remyelination (arrows in **Fig 5J, M**), Some glial processes were present in the lesion (**Fig 5K**, top), and an oligodendrocyte was found in the process of ensheathing an axon (arrow in **Fig 5L**). By contrast, in the cko lesions (**Fig 5 N-Q**), there were many axons without myelin (arrowheads in **Fig 5N**), and the tissue appeared more heterogeneous compared to the control. We could identify numerous glial cells as well as glial processes in the cko tissue (**Fig 5O**). Some had the typical morphological features of oligodendrocytes in the process of myelinating several axons in their proximity (**Fig 5P**).Concomitantly, many axons displayed a glial ensheathment that was not yet compacted (arrows in **Fig 5Q**). These observations suggest that the remyelination process in mg-Nrp1-cko mice was not completely blocked but greatly delayed compared to the controls. Thus, failure to upregulate Nrp1 on activated microglia after acute demyelination significantly impaired the initial proliferative response of OPCs and compromised the timely production of new oligodendrocytes, causing a significant delay in the subsequent remyelination process.

### Exogenous Nrp1 augments PDGF-dependent OPC proliferation by augmenting PDGFRα phosphorylation on OPCs

The above experiments revealed that Nrp1 on microglia was critical for PDGF AA-mediated proliferation of OPCs in the white matter and for the proliferative response of OPCs to acute demyelination. We next examined whether excess Nrp1 could augment the proliferative response of OPCs. Exogenous Nrp1 was added to slice cultures from P8 NG2cre;Z/EG mice in the form of soluble Nrp1-Fc fusion protein. Nrp1-Fc consisted of the extracellular domain of rat Nrp1, which had 98% amino acid identity with mouse Nrp1, fused to the Fc region of human immunoglobulin IgG1 in a homodimeric fusion protein construct. Addition of Nrp1-Fc to slice cultures in the presence of 50 ng/mL PDGF AA led to a significant increase in OPC proliferation in the cortex at 0.4 and 2 μg/mL of Nrp1-Fc (**Fig 6A-C, G**). By contrast, proliferation of OPCs in the corpus callosum was unaffected by all concentrations of Nrp1-Fc tested (**Fig 6D-G**). In the cortex, OPC proliferation reached 36% in the presence of 2 μg/mL Nrp1-Fc but remained lower than the 50% levels seen in white matter.

**Fig 6.**
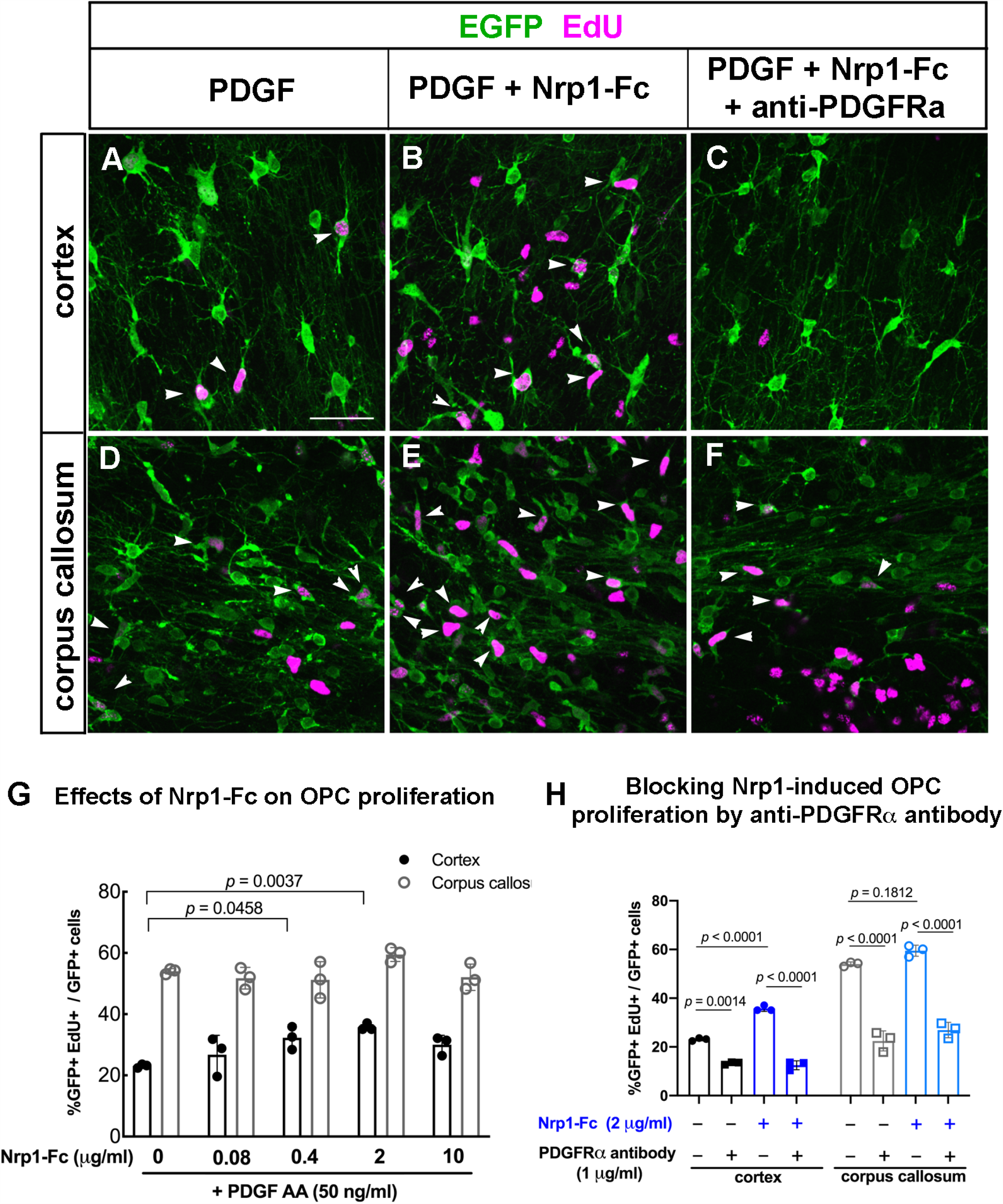
Effects of Nrp1-Fc on OPC proliferation in slice cultures from P8 NG2cre;Z/EG mice. **A-F**. Labeling for EGFP and EdU in the cortex (A-C) and corpus callosum (D-F) of slices treated with 50 ng/mL PDGF AA only (A, D), 50 ng/mL PDGF and 2 μg/mL Nrp1-Fc (B, E), or 50 ng/mL PDGF AA, 2 μg/mL Nrp1-Fc, and 1 μg/mL goat anti-mouse PDGFRα function blocking antibody (C, F). **G**. Scale, 50 μm.Dose-response of OPC proliferation in the cortex and corpus callosum to exogenous Nrp1-Fc in the presence of 50 ng/mL PDGF AA. Values are the percentages of EGFP+ cells that were EdU+. Two-way ANOVA, Tukey’s multiple comparisons test, *n*=3, F(4, 20) = 5.622 for comparisons among different Nrp1-Fc concentrations and F(4, 20) = 317.6 for gray versus white matter comparison. **H**. The effects of anti-PDGFRα blocking antibody on PDGF AA-mediated OPC proliferation in the cortex (gray bars) and corpus callosum (white bars) in the presence (checkered bars) or absence (solid bars) of exogenous Nrp1-Fc. All samples were treated with 50 ng/mL PDGF AA. Two-way ANOVA, Sidak’s multiple comparisons test, *n*=4, F(3, 16) = 266.6.

To determine whether the ability of exogenous Nrp1-Fc to augment PDGF-induced proliferation in gray matter was mediated by PDGFRα, we incubated slice cultures with function-blocking goat antibody to mouse PDGFRα in the presence of PDGF AA. In the absence of Nrp1-Fc, addition of 1 μg/mL of anti-PDGFRα antibody reduced OPC proliferation to 13.3% in gray matter and to 22.5% in white matter (**Fig 6H**). In the presence of Nrp1-Fc, OPC proliferation in the cortex was increased to 35.8%, and this was reduced to 12.5% by anti-PDGFRα antibody (**Fig 6H**). Similarly, addition of anti-PDGFRα antibody to Nrp1-Fc-treated cultures reduced OPC proliferation in white matter to 26.9% (**Fig 6H**), similar to the level in the absence of Nrp1-Fc. These observations indicate that the increased OPC proliferation in gray matter elicited by Nrp1 was mediated through PDGFRα.

Since slice cultures contained different cell types, the above effects of Nrp1-Fc on OPCs could have been mediated by direct effects of Nrp1-Fc on OPCs or by indirectly altering signaling pathways in microglia or other cell types. To resolve this, we added Nrp1-Fc to purified dissociated OPC cultures and examined their proliferation in response to PDGF AA. OPCs were immunopanned from P2-4 mouse neocortex, and 2 μg/mL Nrp1-Fc was added together with 0 to 50 ng/mL PDGF AA (**Fig 7A-E**). Immunopanning yielded an enriched population of OPCs containing <2% F4/80+ microglia. In the presence of PDGF AA concentrations of 4 μg/mL or lower, Nrp1-Fc had no effect on OPC proliferation, and fewer than 4% of Olig2+ OPCs were EdU+ (**Fig 7A-B, E**). In the presence of 50 μg/mL PDGF AA, 54.6% of Olig2+ cells had incorporated EdU without Nrp1-Fc, and this was further increased to 75.9% by the addition of Nrp1-Fc. The most prominent effect of Nrp1-Fc was seen in the presence of 15 μg/mL of PDGF AA, increasing EdU incorporation 3.7-fold from 9% to 33.7% (**Fig 7C-E**). These findings indicate that Nrp1 acted directly on OPCs to augment PDGF AA-induced proliferation, and it was most effective at suboptimum concentrations of PDGF AA.

**Fig 7.**
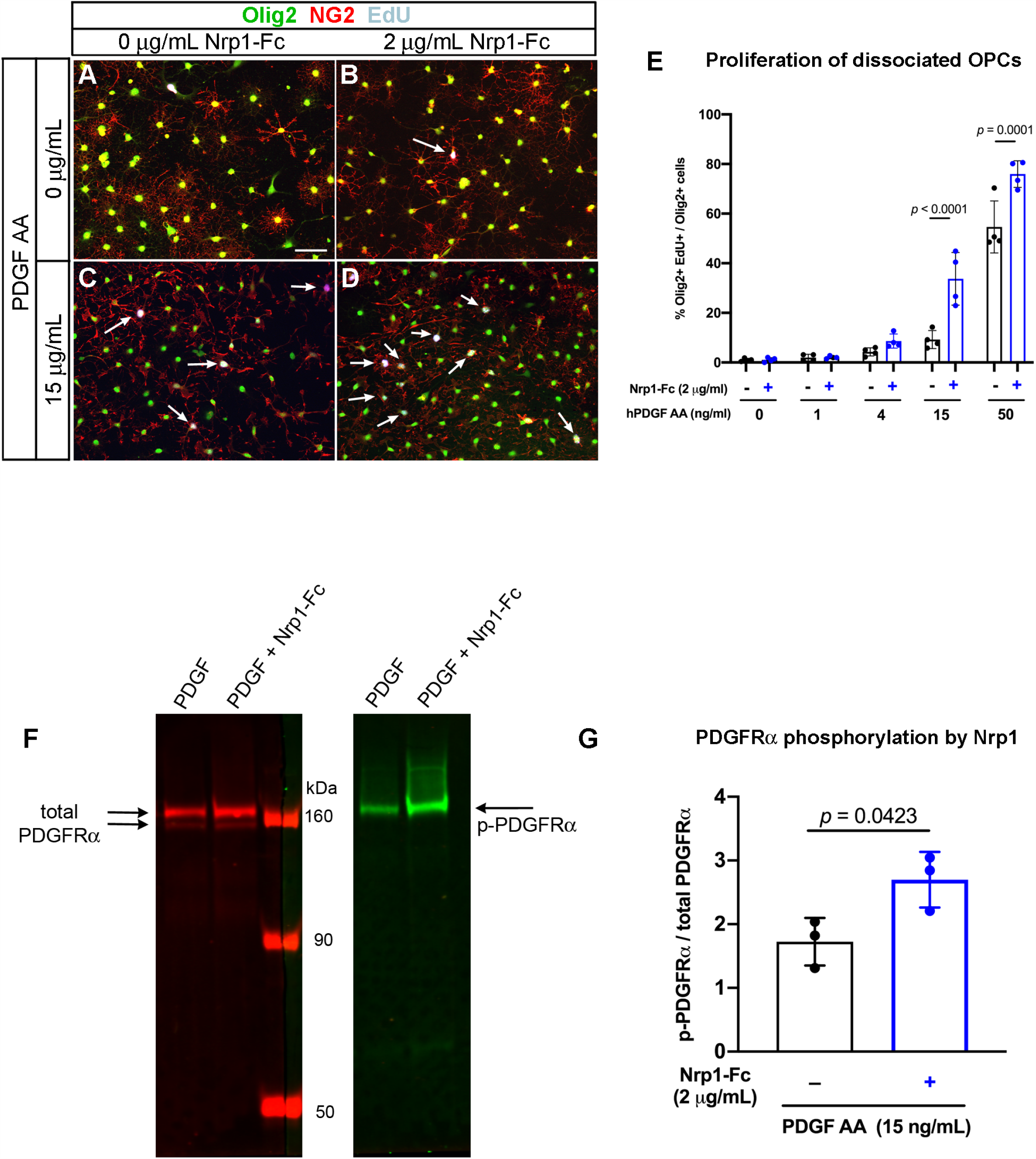
Effects of Nrp1-Fc on dissociated cultures of OPCs. **A-D**. Immunolabeling for Olig2, NG2, and EdU of cells growth in no PDGF AA (A-B) or 15 μg/mL PDGF AA in the absence (A, C) or presence of 2 μg/mL Nrp1-Fc. Scale in A, 50 μm. Arrows: EdU+ Olig2+ NG2+ cells. **E**. Quantification of the effects of combination of PDGF AA and Nrp1-Fc on OPC proliferation. Two-way ANOVA, Sidak’s multiple comparisons test, n=4, F(1, 30) = 36.29. **F**. Immunoblots of OPCs treated with 15 ng/mL of PDGF AA alone or 15 ng/mL of PDGF AA and 2 μg/mL Nrp1-Fc and incubated with antibody to total PDGFRα (left) or Phorphorylated PDGFRα (right). Arrows indicate the expected bands for PDGFRα and phosphorylated PDGFRα. **G**. Quantification of the intensity of the phosphorylated PDGFRα band relative to total PDGFRα bands. ** *p*=0.0081, Student’s paired, two-tailed *t*-test, *n*=3, t = 2.453, df = 2.

Next, we determined whether Nrp1-Fc could augment PDGFRα activation on OPCs exposed to limited amounts of PDGF AA. PDGFRα activation was assessed by immunoblotting for tyrosine phosphorylation on PDGFRα after treating OPCs for 30 minutes at 37°C with 15μg/mL PDGF AA with or without 2 μg/mL Nrp1-Fc. Immunoblots of protein extracts from the treated cells were incubated with rabbit and goat antibodies that recognized phosphorylated PDGFRα and total PDGFRα, respectively. We detected a significant increase in the level of phosphorylated PDGFRα in Nrp1-Fc-treated cells compared with those treated with PDGF AA alone (**Fig 7F-G**). This indicated that Nrp1-Fc could augment the ability of PDGF AA to phosphorylate PDGFRα on OPCs.

To further examine the interaction between Nrp1-Fc and PDGFRα on OPCs, we performed co-clustering experiments by incubating dissociated OPCs with control-Fc or Nrp1-Fc for 30 minutes at 4°C or 36°C and stained for Fc and PDGFRα, as well as Olig2. After incubating OPCs with control-Fc consisting of human IgG Fc dimer at 4°C or 36°C or with Nrp1-Fc at 4°C (**Fig S6 A-C**), PDGFRα was found in small puncta along the processes and some at the soma (A, B), and there was little detectable distinct Fc immunoreactivity. Incubation of OPCs with Nrp1-Fc at 36°C caused greater aggregation of PDGFRα, and Fc immunoreactivity was also found co-clustered with many of the PDGFRα+ aggregates (**Fig S6D**, arrows). These observations further suggest that Nrp1-Fc binds to and causes functional clustering of PDGFRα.

## DISCUSSION

We have shown that Nrp1 expressed by activated microglia plays a critical role in promoting OPC proliferation in early postnatal white matter tracts and after demyelination in the adult corpus callosum. Nrp1 was detected on the majority of amoeboid microglia that appeared transiently in the early postnatal white matter, was not detected on resting ramified microglia, and was re-expressed on activated microglia/macrophages after demyelination. During both development and after demyelination, Nrp1+ microglia were closely apposed to OPC processes. In slice culture, deletion of Nrp1 in microglia significantly reduced PDGF AA-induced OPC proliferation in white but not gray matter. In vivo, microglial Nrp1 deletion significantly reduced OPC proliferation, and the majority of OPCs that were proliferating in P5 corpus callosum were in contact with microglia. Microglial deletion of Nrp1 had no effect in regions where microglia were ramified and did not express Nrp1, such as the cortex and the mature corpus callosum. In adult corpus callosum, Nrp1 was robustly induced on activated microglia/macrophages after acute demyelination, and loss of Nrp1 in these cells significantly reduced OPC proliferation and subsequent oligodendrocyte differentiation and remyelination. Exogenous Nrp1augmented PDGF AA-induced proliferation of cultured OPCs by increasing phosphorylation and clustering of PDGFRα. These findings support the model in which Nrp1 on microglia modulates PDGFRα-mediated proliferation of adjacent OPCs in trans (**Fig 8**).

**Fig 8.**
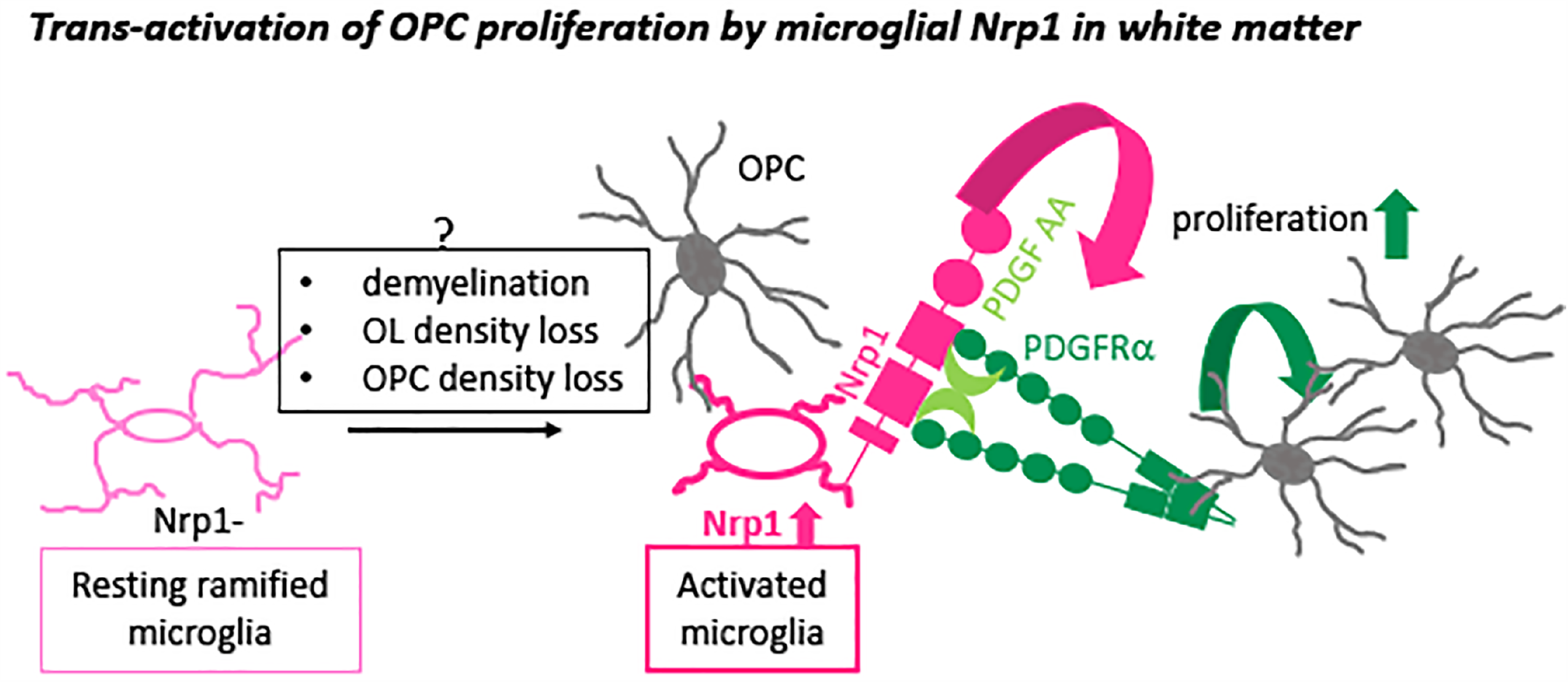
A schematic showing the proposed mechanism of action of microglial Nrp1 on OPC proliferation via activation of PDGFRα in a PDGF AA-dependent manner.

### Dynamic expression of Nrp1 on a subset of “activated” microglia in white matter

CNS parenchymal microglia originate from the embryonic yolk sac and begin to colonize the brain after embryonic day 9.5 (E9.5) ^34^. They exist in different interconvertible functional states. In the normal mature brain, they exist as resting ramified microglia, or “homeostatic microglia”, with highly branched processes ^35,36^ and are involved in a wide range of homeostatic regulation under physiological conditions ^37,38^, as well as in their well known immune function ^39^. In response to a variety of insults, ramified microglia transform into activated phagocytic microglia that become rounded in morphology and upregulate the lysosomal protein CD68 ^40^. Microglia also exist in the early postnatal white matter tracts as amoeboid microglia with phagocytic activity before they transform into ramified microglia ^41^.

Nrp1 was expressed transiently on amoeboid microglia in P5-8 corpus callosum and cerebellar white matter but not on ramified microglia in the cortex and became undetectable on microglia by P30. The developmental window during which Nrp1 was detected on amoeboid microglia correlated with the period during which microglia-specific deletion of Nrp1 reduced OPC proliferation. Furthermore, this coincided with the period of rapid OPC proliferation and OL differentiation.

Resting ramified microglia in the mature CNS no longer expressed Nrp1. However, within three days following acute demyelination, activated CD68+ microglia/macrophages robustly upregulated Nrp1 on the cell surface. Deletion of Nrp1 from microglia significantly reduced OPC proliferation in the lesion during the first week after demyelination and subsequently led to reduced OL differentiation and myelination. Although the signals that dynamically upregulate Nrp1 on activated microglia in the developing and demyelinated white matter remain unknown, it is possible that phagocytosis of dead OL lineage cells by amoeboid/activated microglia, a process which has been previously reported ^41,42^, could trigger a response that activates their Nrp1 expression.

#### Transcriptomic similarities between early postnatal amoeboid microglia and demyelination-induced activated microglia

Recent single-cell transcriptomic studies have identified a transcriptionally unique subtype of microglia/monocytic cells in the mouse brain that are found in early postnatal white matter, which are referred to as axon tract-associated microglia (ATM) ^43^ or early postnatal proliferative region-associated microglia (PAM) ^44^, where oligodendrogliogenesis is actively occurring prior to myelination. Intriguingly, among microglia from five different ages and from demyelinated lesions, Nrp1 transcript counts were highest in those from P4/5 brain and LPC-induced white matter lesions ^43^http://www.microgliasinglecell.com/), whereas Nrp1 expression was lower in the homeostatic microglia in the mature CNS. Thus, the two subpopulations of phagocytic microglia not only share their transcriptome but they also regulate OPC proliferation similarly by dynamically modulating Nrp1 expression on their surface.

### Beneficial effects of microglia on OL/myelin production

Traditionally, microglia have been associated with inflammation and deleterious effects on OLs and myelin in demyelinating lesions of multiple sclerosis 45. However, beneficial effects of microglia during myelin repair have also been identified ^46^. Their dichotomous role appears to depend on their activation state ^47,48^. During normal development, amoeboid microglia/ATM/PAM in the early postnatal white matter tracts play a critical role in myelination. This is mediated in part by insulin-like growth factor-1 (IGF1) ^49^, which is highly expressed in amoeboid microglia ^43,44^. Pharmacological or genetic perturbation of the activation state of the amoeboid microglia causes deficits in OL development and myelination ^50-52^. However, the mechanisms by which microglia impart these positive effects on OL lineage cells have remained unclear.

Our studies have revealed that Nrp1, which is highly expressed on activated microglia in white matter, plays a critical role in promoting OPC proliferation during development and after demyelination. Developmental deletion of microglial Nrp1 did not lead to permanent myelin defects, which could be due to compensation by proliferation and migration of microglia that had escaped Nrp1 deletion. Since we only deleted Nrp1 from microglia and not from vascular cells or axons, the partial inhibition of OPC proliferation in the mg-Nrp1-cko could be due to additional effects of Nrp1 on cell types other than microglia, as well as Nrp1-independent mechanisms that also promote developmental OPC proliferation.

Nrp1 loss from activated microglia that appeared in the demyelinated lesion led to a severe decrease in OPC proliferation followed by decreased OL regeneration and protracted remyelination. With compromised proliferative response of OPCs after acute demyelination, additional OPCs may have to be recruited from surrounding areas such as the subventricular zone or the gray matter, which could have contributed to the delayed remyelination. Interestingly, Nrp1 is highly expressed by microglia/macrophages associated with malignant glioma ^53^, and deletion of Nrp1 from microglia and macrophages slows glioma growth ^54^.

### Mechanism by which microglial Nrp1 promotes OPC proliferation

Nrp1 is the axonal co-receptor for class III Semaphorins ^21,55^ and is critical for growth cone collapse induced by Sema3A ^56^. Nrp1 was highly expressed on axons in E18.5 corpus callosum but the level had significantly dropped by P5. Axonal Nrp1 became undetectable in the corpus callosum after P14 and was not re-expressed after demyelination. Thus, axonal Nrp1 is not likely to be a major modulator of OPC proliferation. A related protein Nrp2 shares 44% amino acid identity with Nrp1 and binds primarily Sema3F. Semaphorins have been implicated in OPC migration during development ^57,58^ and after demyelination, acting via Nrp1 and 2 on OPCs ^59,60^. However, we did not detect Nrp1 or 2 expression on OPCs, or on other cells in the corpus callosum, which was consistent with RNA-seq data for OPCs ^61,62^. Neither did we detect Nrp2 on microglia, also consistent with the transcriptomic studies ^43,44,61^. Furthermore, the dramatic decrease in OPC proliferation in demyelinated lesions after Nrp1 deletion from activated microglia supports the view that microglial Nrp1 functions as a key inducer of OPC proliferation.

#### Different saturation levels of Nrp1 in gray and white matter differentially affect OPC proliferation

In slice cultures, blocking Nrp1 reduced PDGF AA-mediated OPC proliferation in the corpus callosum but not in the cortex. By contrast, addition of Nrp1-Fc augmented PDGF AA-mediated OPC proliferation in gray but not white matter. Nrp1 may be present in saturating amounts in white matter, whereas in gray matter, the level of endogenous Nrp1 may be insufficient to adequately activate PDGFRα on OPCs. This is consistent with the immunohistochemical detection of higher levels of Nrp1 on activated microglia in white matter. This is also supported by the ability of Nrp1-Fc to augment proliferation and PDGFRα phosphorylation on dissociated OPCs in the presence of a suboptimum concentration of PDGF AA.

#### Trans-activation of PDGFRα on OPCs by Nrp1 on adjacent microglia

On mesenchymal stem cells, Nrp1 forms a complex with PDGFRα in the presence of PDGF and increases their migration and proliferation in response to PDGF AA ^18^. In this case, both Nrp1 and PDGFRα are expressed in cis on the same cell, similar to the role of Nrp1 in VEGF_165_-mediated VEGFR2 stimulation (reviewed in ^22,63^). By contrast, our findings support a model of trans-activation of PDGFRα on OPCs by Nrp1 on adjacent microglia (**Fig 8**). Juxtacrine interaction between NRP1-expressing tumor cells and VEGFR2-endothelial cells can occur in vitro in the presence of VEGF_165_ ^64^. In porcine aortic endothelial cells, presentation of human NRP1 in trans to VEGFR2 reduces internalization of VEGFR2 on the cell surface and causes delayed and sustained intracellular effects compared to cis-activation of VEGFR2 by NRP1 ^65^.Our observation that Nrp1-Fc co-clustered with PDGFRα on cultured OPCs and enhanced its phosphorylation is also consistent with the role of Nrp1 in stabilizing PDGFRα on the OPC surface. It is also possible that Nrp1 increases the availability of PDGF AA to PDGFRα by forming a ternary complex.

On other cell types, Nrp1 has been shown to be released by extracellular proteases ^66,67^. However, if Nrp1 released from microglia were to affect OPC proliferation, the diffusion radius would have to be limited to the immediate pericellular environment, as the effects were confined to the corpus callosum and did not extend to the cortex. Furthermore, the immunofluorescence signal for Nrp1 was most prominent along the surface of microglia (**Fig 4D and Fig S2**). We previously reported a strikingly close apposition between an OPC and its adjacent microglia ^68^. These observations, along with the close apposition between Nrp1+ microglia and OPCs observed here (**Fig 2**), support a close contact-mediated interaction in trans. It is likely that Nrp1 signal constitutes a part of a larger constellation of microglia-to-OPC/OL signaling that maintains the homeostasis of OL and myelin density. Future studies may be directed to exploring the signaling mechanism of the cross-talk between Nrp1-expressing microglia and OPCs and how such interactionscontribute to OL/myelin homeostasis.

## ACKNOWLEDGMENTS

We thank Youfen Sun for maintaining the mouse colony. This study was supported by NIH R01 NS073425 (AN) and funding from the European Union’s Framework Program for Research and Innovation Horizon 2020(2014-2020) under the Marie Sklodowska-Curie Grant Agreement No. 845366 (FP). We thank Dr. Chris O’Connell, Director of Advanced Light Microscopy Core, for his assistance with the Leica SP8 confocal microscope, which was purchased with funds from an NIH Instrumentation Grant S10 OD016435 (PI: Akiko Nishiyama). Electron microscopy was performed at the Biosciences Electron Microscopy Facility of the University of Connecticut under the guidance of Director Dr. Linnaea Ostroff and with the excellent technical support of Drs. Maritza Abril and Xuanhao Sun. We thank Dr. Timothy Moore, Director of the Statistical Consulting Services, for his assistance with statistical analyses and graphing of the data.

## Author contributions

AS, AR, FP, and AN designed the study. AS, AR, FP, and WMW generated data. AS, FP, WMW, and AN interpreted the data. AS, FP, and AN wrote the manuscript, and AS, FP, WMW, and AN edited the manuscript.

## Declaration of competing interests

The authors have no competing interests to declare.

## Data availability statement

The datasets generated and analyzed in this study are available from the corresponding author upon reasonable request.

## FIGURE LEGENDS

**Supplemental Fig 1**. The distribution of Nrp1 in the developing CNS

**A**. Nrp1 expression on laminin+ blood vessels in P5 cortex. Scale, 20 μm.

**B**. P5 thoracic spinal cord from Cx3CR1^creERT2-ires-EYFP^ mouse stained for Nrp1. YFP+ microglia in the white matter are Nrp1-negative (arrows). Strong neuronal Nrp1 expression is seen in the dorsal spinal cord. Scale, 50 μm.

**C**. Coronal section of forebrain from E18.5 Cx3CR1^creERT2-ires-EYFP^ mouse showing Nrp1 on axons in the dorsal corpus callosum. YFP+ microglia in the corpus callosum do not express Nrp1 (arrowheads), while the round YFP+ meningeal macrophage-like cells near the septum are Nrp1+ (arrows). Scale, 50 μm. ctx, cortex; cc, corpus callosum.

**Supplemental Fig 2**. Nrp1 expression on EYFP+ microglia in the cortex and corpus callosum during postnatal development of Cx3CR1^creERT2-ires-EYFP^ mice. Arrows, examples of blood vessels. Arrowheads, examples of Nrp1+ EYFP+ microglia. Scale bar in A = 50 μm.

**Supplemental Fig 3. CD68 is expressed on Nrp1+ amoeboid microglia**.

**A-D**. CD68 is detected in some Nrp1+ amoeboid microglia in P5 corpus callosum (arrows). Scale, 50 μm.

**E-H**. In P14 corpus callosum, Nrp1 is no longer detected on microglia that appear ramified. These microglia have significantly less CD68 immunoreactivity than those in P5 corpus callosum, and it is restricted a few puncta (arrowheads). Scale, 50 μm.

**Supplemental Fig 4. Characterization of mg-Nrp1-cko**.

**A**. Schematic of generating microglia-specific Nrp1 knock out (mg-Nrp1-cko) and the heterozygous control (mg-Nrp1-cont).

**B-C**. P5 corpus callosum after 4OHT injection at P2-3, showing a complete absence of Nrp1 on EYFP+ microglia (arrowheads show examples) but retained on blood vessels (arrow). Scale, 50μm.

**D**. The proportion of Nrp1+ EYFP+ cells among EYFP+ microglia in the corpus callosum of P5 mg-Nrp1-cont and mg-Nrp1-cko mice. Student’s t-test, n = 3, t = 156.9, df =4.

**E**. The density of Nrp1+ EYPF+ microglia in the corpus callosum of P5 mg-Nrp1-cont and mg-Nrp1-cko mice. Student’s t-test, n = 3, t = 0.7434, df =4.

**F**. The density of Aldh1L1+ astrocytes in the corpus callosum of P5 mg-Nrp1-cont and mg-Nrp1-cko mice. Student’s t-test, n = 3, t = 0.3116, df =4.

**G** The proportion of the corpus callosum area occupied by blood vessels outlined by laminin immunoreactivity. Student’s t-test, n = 3, t = 0.5039, df =4.

**H-J**. TUNEL labeling of P5 corpus callosum of mg-Nrp1-cont (**F**) and mg-Nrp1-cko (**G**) and the external granule cell layer of the cerebellar cortex of mg-Nrp1-cont, showing a positive cell (**H**). Scale 20 μm.

**K-P**. Immunolabeling for MBP in the corpus callosum of P14 mg-Nrp1-cont and cko mice. Scale 50 μm.

**Supplemental Fig 5**. PDGF AA-mediated OPC proliferation in mg-Nrp1-cont and cko.

**B**. Schematic showing EdU labeling of slice cultures from P8 mg-Nrp1-cont or cko mice after cre induction in vivo at P2 and P3.

**C**. Quantification showing the proportion of NG2+ cells that were EdU+ in PDGF AA-treated slices. Two-way ANOVA, Tukey’s multiple comparisons test, *n*=3, F(1, 8) = 100.4.

**Supplemental Fig 6**. Co-clustering of Nrp1-Fc with PDGFRα on dissociated OPCs.

**A-B**. Immunopanned OPCs were incubated for 30 minutes with control human IgG Fc dimer (Cont-Fc) at 4°C (A) or 36°C (B) and stained for PDGFRα and Fc.

**C-D**. Immunopanned OPCs were incubated for 30 minutes with Nrp1-Fc fusion protein (Nrp1-Fc) at 4°C (A) or 36°C (B) and stained for PDGFRα and Fc.

Arrows show Nrp1-Fc co-clustered with PDGFRα. Scale, 20 μm.

## Notes

### Competing Interest Statement

The authors have declared no competing interest.

## REFERENCES

Dawson, M.R., Polito, A., Levine, J.M. & Reynolds, R. NG2-expressing glial progenitor cells: an abundant and widespread population of cycling cells in the adult rat CNS. Mol Cell Neurosci 24, 476–88 (2003).

Nishiyama, A., Komitova, M., Suzuki, R. & Zhu, X. Polydendrocytes (NG2 cells): multifunctional cells with lineage plasticity. Nat Rev Neurosci 10, 9–22 (2009).

Nishiyama, A., Boshans, L., Goncalves, C.M., Wegrzyn, J. & Patel, K.D. Lineage, fate, and fate potential of NG2-glia. Brain Res 1638, 116–28 (2016).

Richardson, W.D., Pringle, N., Mosley, M.J., Westermark, B. & Dubois-Dalcq, M. A role for platelet-derived growth factor in normal gliogenesis in the central nervous system. Cell 53, 309–319 (1988).

Calver, A.R. et al. Oligodendrocyte population dynamics and the role of PDGF in vivo. Neuron 20, 869–882 (1998).

Fruttiger, M., Calver, A.R. & Richardson, W.D. Platelet-derived growth factor is constitutively secreted from neuronal cell bodies but not from axons. Curr Biol 10, 1283–6. (2000).

Wu, Q. et al. Elevated levels of the chemokine GRO-1 correlate with elevated oligodendrocyte progenitor proliferation in the jimpy mutant. Journal of Neuroscience 20, 2609–2617 (2000).

Bu, J., Banki, A., Wu, Q. & Nishiyama, A. Increased NG2(+) glial cell proliferation and oligodendrocyte generation in the hypomyelinating mutant shiverer. Glia 48, 51–63 (2004).

Gensert, J.M. & Goldman, J.E. Endogenous progenitors remyelinate demyelinated axons in the adult CNS. Neuron 19, 197–203 (1997).

Keirstead, H.S., Levine, J.M. & Blakemore, W.F. Response of the oligodendrocyte progenitor cell population (defined by NG2 labelling) to demyelination of the adult spinal cord. Glia 22, 161–170 (1998).

Di Bello, C.I., Dawson, M.R., Levine, J.M. & Reynolds, R. Generation of oligodendroglial progenitors in acute inflammatory demyelinating lesions of the rat brain stem is associated with demyelination rather than inflammation. J Neurocytol 28, 365–81. (1999).

Watanabe, M., Toyama, Y. & Nishiyama, A. Differentiation of proliferated NG2-positive glial progenitor cells in a remyelinating lesion. J Neurosci Res 69, 826–36. (2002).

Hill, R.A. & Nishiyama, A. NG2 cells (polydendrocytes): listeners to the neural network with diverse properties. Glia 62, 1195–210 (2014).

Boshans, L.L., Sherafat, A. & Nishiyama, A. The effects of developmental and current niches on oligodendrocyte precursor dynamics and fate. Neurosci Lett (2020).

Hill, R.A., Patel, K.D., Medved, J., Reiss, A.M. & Nishiyama, A. NG2 cells in white matter but not gray matter proliferate in response to PDGF. J Neurosci 33, 14558–66 (2013).

Vigano, F., Mobius, W., Gotz, M. & Dimou, L. Transplantation reveals regional differences in oligodendrocyte differentiation in the adult brain. Nat Neurosci 16, 1370–2 (2013).

Andrae, J., Gallini, R. & Betsholtz, C. Role of platelet-derived growth factors in physiology and medicine. Genes Dev 22, 1276–312 (2008).

Ball, S.G., Bayley, C., Shuttleworth, C.A. & Kielty, C.M. Neuropilin-1 regulates platelet-derived growth factor receptor signalling in mesenchymal stem cells. Biochem J 427, 29–40 (2010).

Takagi, S. et al. The A5 antigen, a candidate for the neuronal recognition molecule, has homologies to complement components and coagulation factors. Neuron 7, 295–307 (1991).

Kawakami, A., Kitsukawa, T., Takagi, S. & Fujisawa, H. Developmentally regulated expression of a cell surface protein, neuropilin, in the mouse nervous system. J Neurobiol 29, 1–17 (1996).

He, Z. & Tessier-Lavigne, M. Neuropilin is a receptor for the axonal chemorepellent Semaphorin III. Cell 90, 739–51 (1997).

Zachary, I.C. How neuropilin-1 regulates receptor tyrosine kinase signalling: the knowns and known unknowns. Biochem Soc Trans 39, 1583–91 (2011).

Novak, A., Guo, C., Yang, W., Nagy, A. & Lobe, C.G. Z/EG, a double reporter mouse line that expresses enhanced green fluorescent protein upon Cre-mediated excision. Genesis 28, 147–55. (2000).

Zhu, X. et al. Age-dependent fate and lineage restriction of single NG2 cells. Development 138, 745–53 (2011).

Zhu, X., Bergles, D.E. & Nishiyama, A. NG2 cells generate both oligodendrocytes and gray matter astrocytes. Development 135, 145–57 (2008).

Gu, C. et al. Neuropilin-1 conveys semaphorin and VEGF signaling during neural and cardiovascular development. Dev Cell 5, 45–57 (2003).

Parkhurst, C.N. et al. Microglia promote learning-dependent synapse formation through brain-derived neurotrophic factor. Cell 155, 1596–609 (2013).

Serwanski, D.R., Rasmussen, A.L., Brunquell, C.B., Perkins, S.S. & Nishiyama, A. Sequential Contribution of Parenchymal and Neural Stem Cell-Derived Oligodendrocyte Precursor Cells toward Remyelination. Neuroglia 1, 91–105 (2018).

Sherafat, A., Hill, R.A. & Nishiyama, A. Organotypic Slice Cultures to Study Oligodendrocyte Proliferation, Fate, and Myelination. Methods Mol Biol 1791, 145–156 (2018).

Dugas, J.C. & Emery, B. Purification and culture of oligodendrocyte lineage cells. Cold Spring Harb Protoc 2013, 810–4 (2013).

Austyn, J.M. & Gordon, S. F4/80, a monoclonal antibody directed specifically against the mouse macrophage. Eur.J.Immunol. 11, 805–815 (1981).

Holness, C.L. & Simmons, D.L. Molecular cloning of CD68, a human macrophage marker related to lysosomal glycoproteins. Blood 81, 1607–13 (1993).

Hughes, M.M., Field, R.H., Perry, V.H., Murray, C.L. & Cunningham, C. Microglia in the degenerating brain are capable of phagocytosis of beads and of apoptotic cells, but do not efficiently remove PrPSc, even upon LPS stimulation. Glia 58, 2017–30 (2010).

Ginhoux, F. et al. Fate mapping analysis reveals that adult microglia derive from primitive macrophages. Science 330, 841–5 (2010).

Perry, V.H. & Gordon, S. Macrophages and microglia in the nervous system. Trends in the Neurosciences 11, 273–277 (1988).

Kettenmann, H., Hanisch, U.K., Noda, M. & Verkhratsky, A. Physiology of microglia. Physiol Rev 91, 461–553 (2011).

Schafer, D.P. & Stevens, B. Microglia Function in Central Nervous System Development and Plasticity. Cold Spring Harb Perspect Biol 7, a020545 (2015).

Butovsky, O. & Weiner, H.L. Microglial signatures and their role in health and disease. Nat Rev Neurosci 19, 622–635 (2018).

Owens, T. Immune Functions of Microglia. in Neuroglia (ed. Ransom, H.K.a.B.R.) 638–648 (Oxford University Press, New York, 2013).

Streit, W.J. Microglial Cells. in Neuroglia (ed. Ransom, H.K.a.B.R.) 86-97 (Oxford University Press, New York, 2013).

Ling, E.A. Some aspects of amoeboid microglia in the corpus callosum and neighbouring regions of neonatal rats. J Anat 121, 29–45 (1976).

Ellison, J.A. & de Vellis, J. Amoeboid microglia expressing G<sub>D3</sub> ganglioside are concentrated in regions of oligodendrogenesis during development of the rat corpus callosum. Glia 14, 123–132 (1995).

Hammond, T.R. et al. Single-Cell RNA Sequencing of Microglia throughout the Mouse Lifespan and in the Injured Brain Reveals Complex Cell-State Changes. Immunity 50, 253– 271 e6 (2019).

Li, Q. et al. Developmental Heterogeneity of Microglia and Brain Myeloid Cells Revealed by Deep Single-Cell RNA Sequencing. Neuron 101, 207–223 e10 (2019).

Lassmann, H. Multiple sclerosis: lessons from molecular neuropathology. Exp Neurol 262 Pt A, 2–7 (2014).

Miron, V.E. et al. M2 microglia and macrophages drive oligodendrocyte differentiation during CNS remyelination. Nat Neurosci 16, 1211–8 (2013).

Nicholas, R.S., Wing, M.G. & Compston, A. Nonactivated microglia promote oligodendrocyte precursor survival and maturation through the transcription factor NF-kappa B. Eur J Neurosci 13, 959–67 (2001).

Butovsky, O. et al. Induction and blockage of oligodendrogenesis by differently activated microglia in an animal model of multiple sclerosis. J Clin Invest 116, 905–15 (2006).

Wlodarczyk, A. et al. A novel microglial subset plays a key role in myelinogenesis in developing brain. EMBO J 36, 3292–3308 (2017).

Hagemeyer, N. et al. Microglia contribute to normal myelinogenesis and to oligodendrocyte progenitor maintenance during adulthood. Acta Neuropathol 134, 441–458 (2017).

Erblich, B., Zhu, L., Etgen, A.M., Dobrenis, K. & Pollard, J.W. Absence of colony stimulation factor-1 receptor results in loss of microglia, disrupted brain development and olfactory deficits. PLoS One 6, e26317 (2011).

Shigemoto-Mogami, Y., Hoshikawa, K., Goldman, J.E., Sekino, Y. & Sato, K. Microglia enhance neurogenesis and oligodendrogenesis in the early postnatal subventricular zone. J Neurosci 34, 2231–43 (2014).

Caponegro, M.D., Moffitt, R.A. & Tsirka, S.E. Expression of neuropilin-1 is linked to glioma associated microglia and macrophages and correlates with unfavorable prognosis in high grade gliomas. Oncotarget 9, 35655–35665 (2018).

Miyauchi, J.T. et al. Deletion of Neuropilin 1 from Microglia or Bone Marrow-Derived Macrophages Slows Glioma Progression. Cancer Res 78, 685–694 (2018).

Kolodkin, A.L. et al. Neuropilin is a semaphorin III receptor. Cell 90, 753–62 (1997).

Luo, Y., Raible, D. & Raper, J.A. Collapsin: a protein in brain that induces the collapse and paralysis of neuronal growth cones. Cell 75, 217–27 (1993).

Sugimoto, Y. et al. Guidance of glial precursor cell migration by secreted cues in the developing optic nerve. Development 128, 3321–30. (2001).

Spassky, N. et al. Directional guidance of oligodendroglial migration by class 3 semaphorins and netrin-1. J Neurosci 22, 5992–6004 (2002).

Williams, A. et al. Semaphorin 3A and 3F: key players in myelin repair in multiple sclerosis? Brain 130, 2554–65 (2007).

Piaton, G. et al. Class 3 semaphorins influence oligodendrocyte precursor recruitment and remyelination in adult central nervous system. Brain 134, 1156–67 (2011).

Zhang, Y. et al. An RNA-sequencing transcriptome and splicing database of glia, neurons, and vascular cells of the cerebral cortex. J Neurosci 34, 11929–47 (2014).

Marques, S. et al. Transcriptional Convergence of Oligodendrocyte Lineage Progenitors during Development. Dev Cell 46, 504–517 e7 (2018).

Pellet-Many, C., Frankel, P., Jia, H. & Zachary, I. Neuropilins: structure, function and role in disease. Biochem J 411, 211–26 (2008).

Soker, S., Miao, H.Q., Nomi, M., Takashima, S. & Klagsbrun, M. VEGF165 mediates formation of complexes containing VEGFR-2 and neuropilin-1 that enhance VEGF165-receptor binding. J Cell Biochem 85, 357–68 (2002).

Koch, S. et al. NRP1 presented in trans to the endothelium arrests VEGFR2 endocytosis, preventing angiogenic signaling and tumor initiation. Dev Cell 28, 633–46 (2014).

Mehta, V. et al. VEGF (Vascular Endothelial Growth Factor) Induces NRP1 (Neuropilin-1) Cleavage via ADAMs (a Disintegrin and Metalloproteinase) 9 and 10 to Generate Novel Carboxy-Terminal NRP1 Fragments That Regulate Angiogenic Signaling. Arterioscler Thromb Vasc Biol 38, 1845–1858 (2018).

Xu, D. et al. Novel MMP-9 substrates in cancer cells revealed by a label-free quantitative proteomics approach. Mol Cell Proteomics 7, 2215–28 (2008).

Nishiyama, A., Yu, M., Drazba, J.A. & Tuohy, V.K. Normal and reactive NG2+ glial cells are distinct from resting and activated microglia. Journal of Neuroscience Research 48, 299–312 (1997).

