## Supplemental Figure 1-6 for "Microglial Neuropilin-1 trans-regulates oligodendrocyte expansion during development and remyelination"

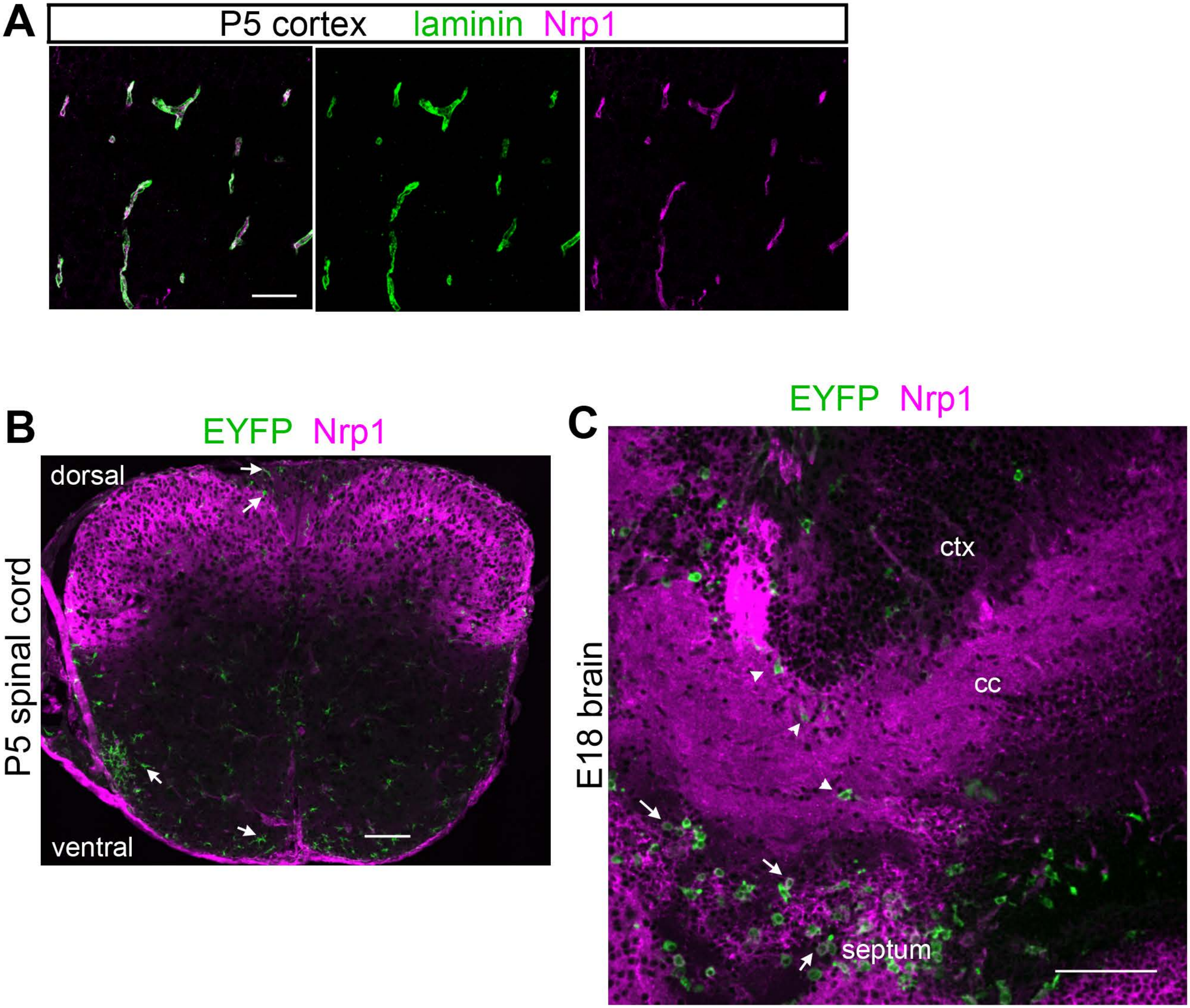


Sherafat et al., Supplemental Figure 1

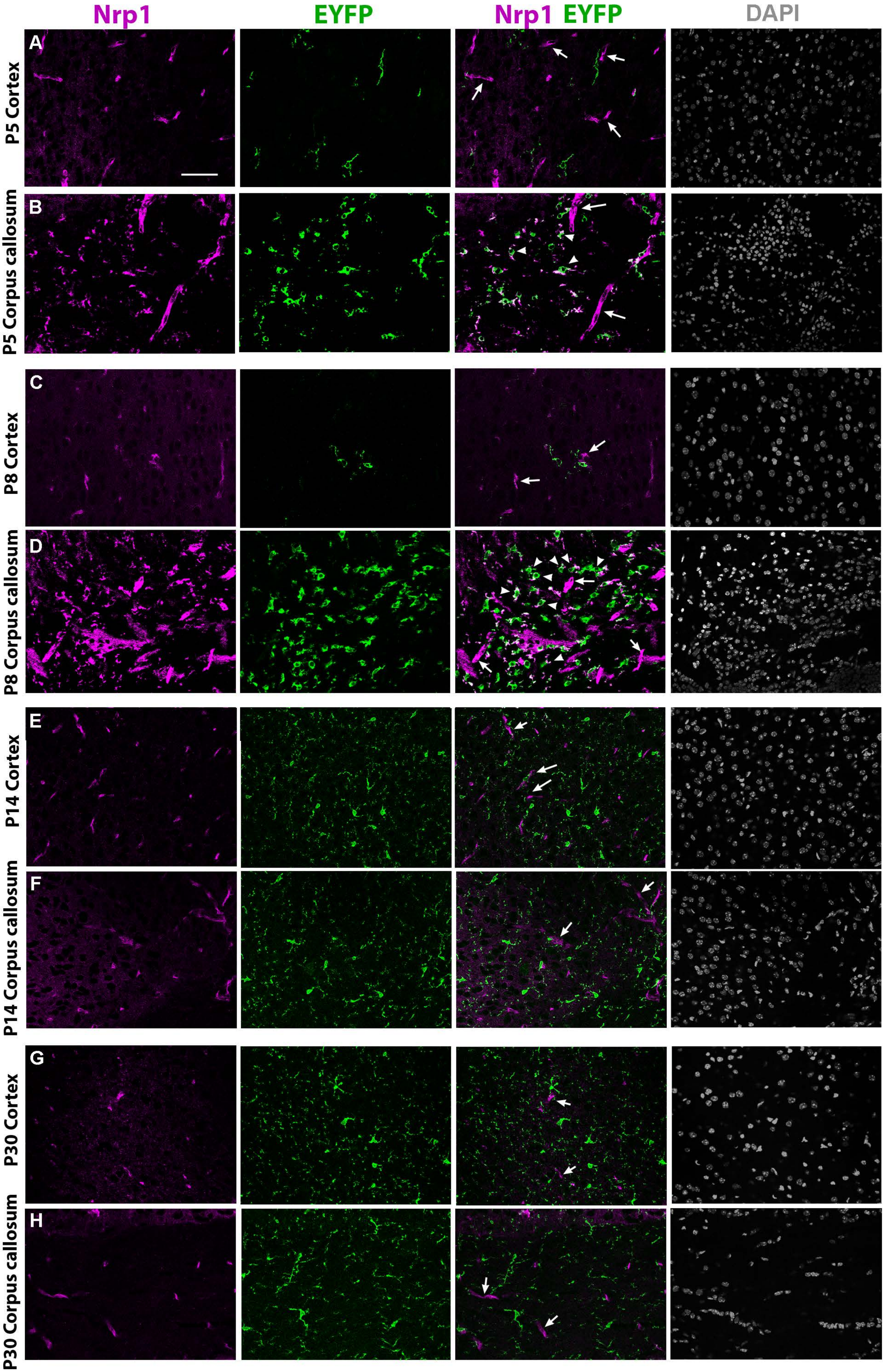


Sherafat et al., Supplemental Figure 2

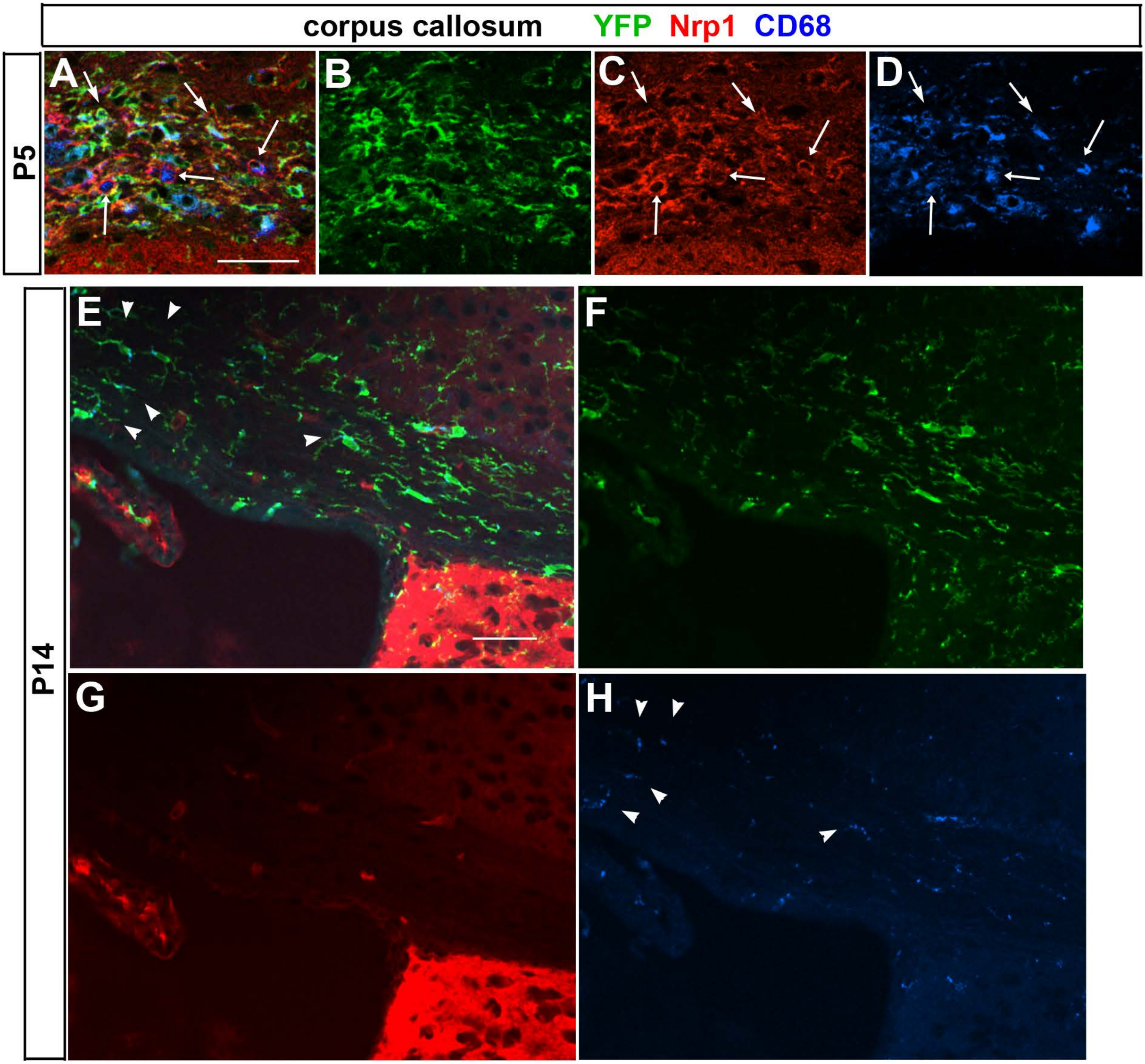


Sherafat et al., Figure S3

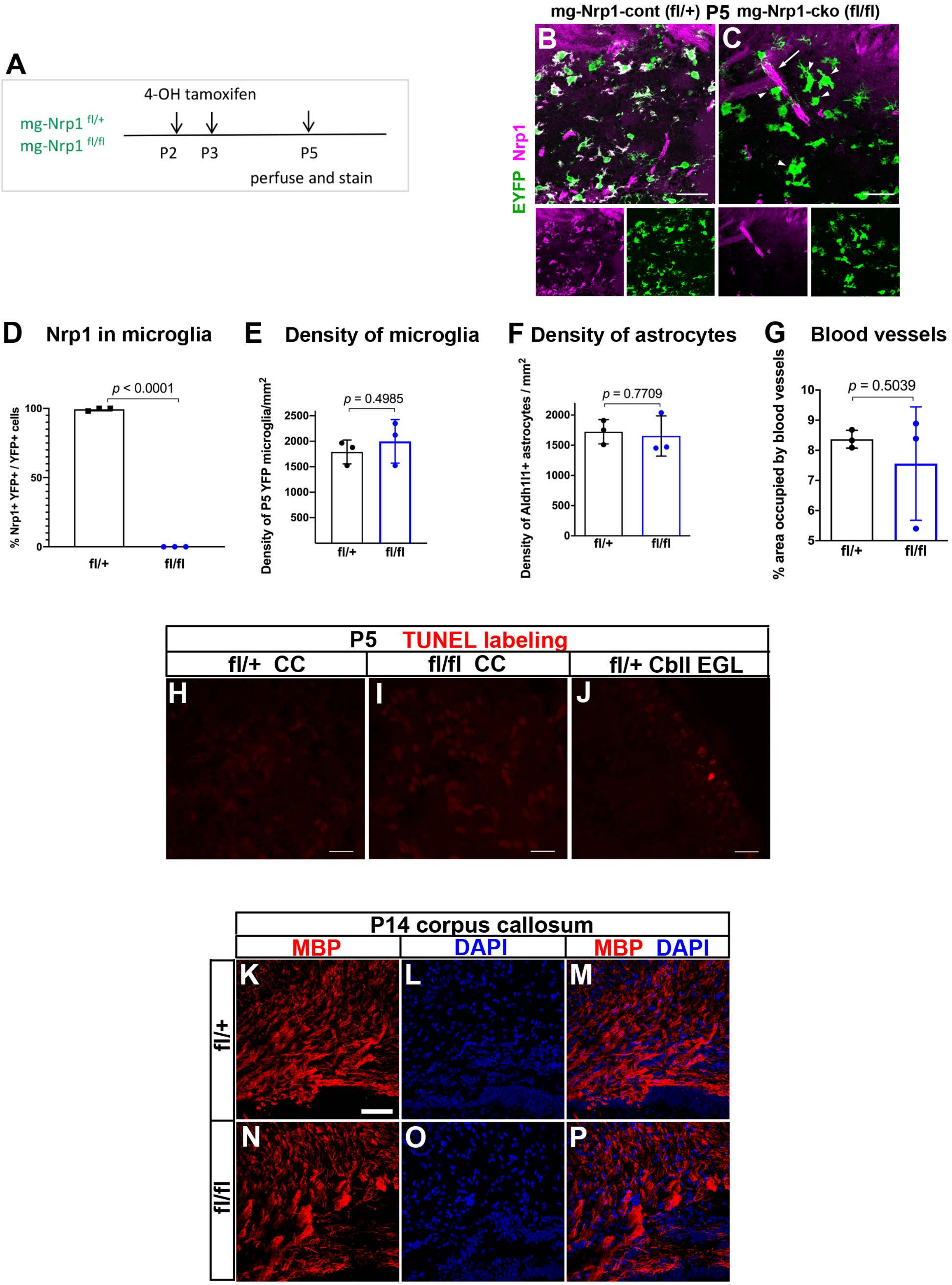


Sherafat et al., Supplemental Figure 4

A

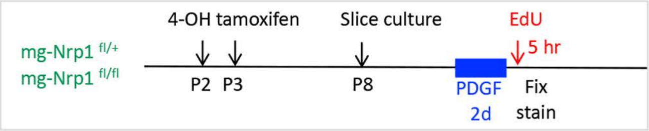


**PDGF AA-dependent OPC proliferation in slice culture**

B

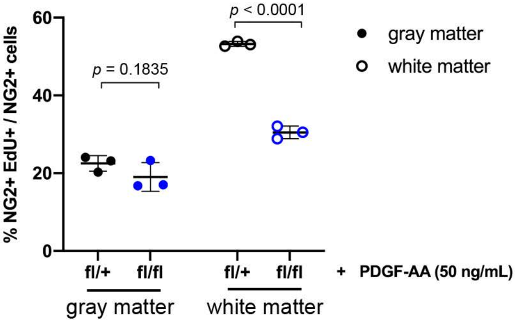


Sherafat et al., Supplemental Figure 5

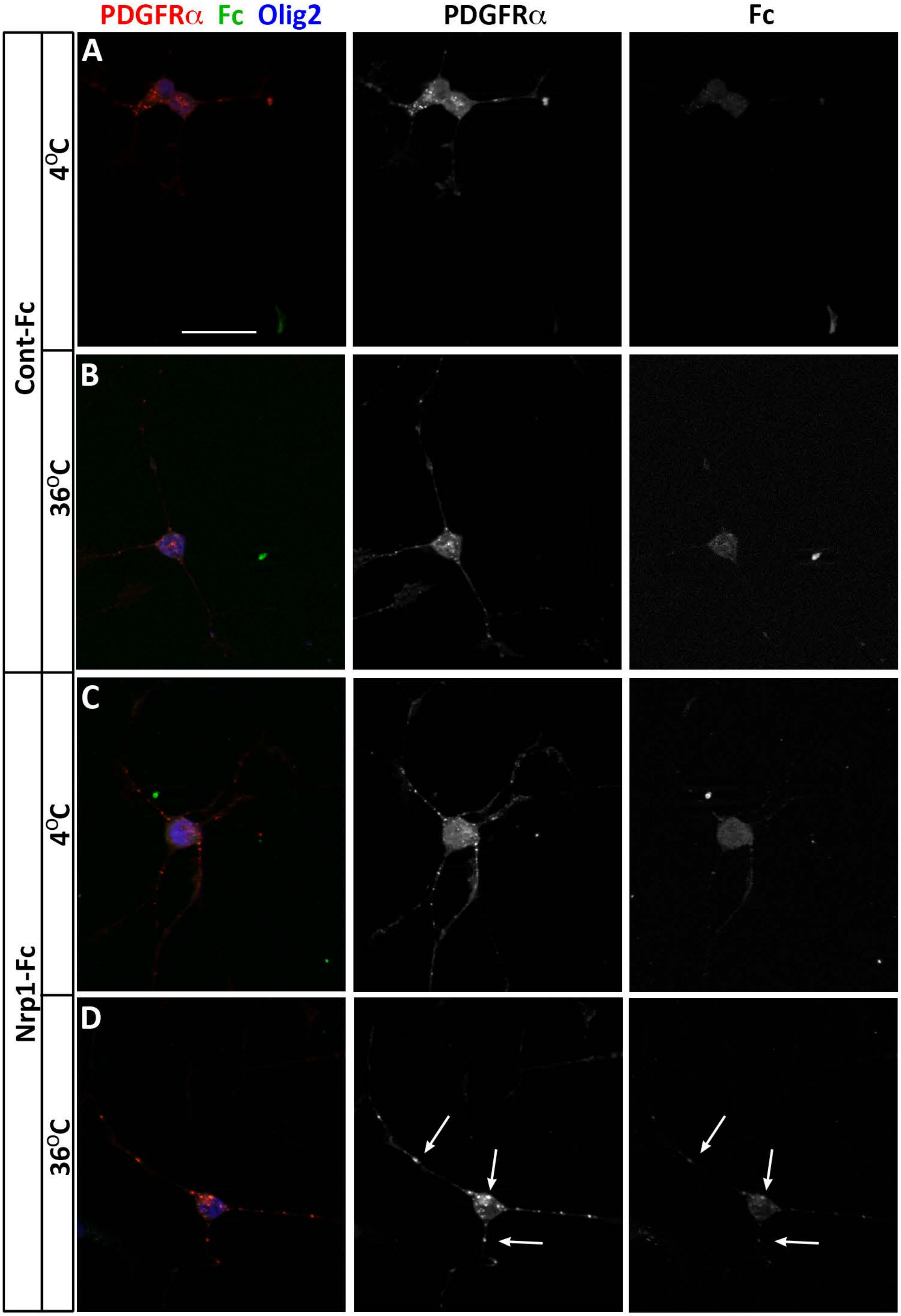


Sherafat et al., Supplemental Figure 6
